# Robust phenotyping of highly multiplexed tissue imaging data using pixel-level clustering

**DOI:** 10.1101/2022.08.16.504171

**Authors:** Candace C. Liu, Noah F. Greenwald, Alex Kong, Erin F. McCaffrey, Ke Xuan Leow, Dunja Mrdjen, Bryan J. Cannon, Josef Lorenz Rumberger, Sricharan Reddy Varra, Michael Angelo

## Abstract

While technologies for multiplexed imaging have provided an unprecedented understanding of tissue composition in health and disease, interpreting this data remains a significant computational challenge. To understand the spatial organization of tissue and how it relates to disease processes, imaging studies typically focus on cell-level phenotypes. However, images can capture biologically important objects that are outside of cells, such as the extracellular matrix. Here, we developed a pipeline, Pixie, that achieves robust and quantitative annotation of pixel-level features using unsupervised clustering and show its application across a variety of biological contexts and multiplexed imaging platforms. Furthermore, current cell phenotyping strategies that rely on unsupervised clustering can be labor intensive and require large amounts of manual cluster adjustments. We demonstrate how pixel clusters that lie within cells can be used to improve cell annotations. We comprehensively evaluate pre-processing steps and parameter choices to optimize clustering performance and quantify the reproducibility of our method. Importantly, Pixie is open source and easily customizable through a user-friendly interface.

## Introduction

The advancement of multiplexed tissue imaging technologies over the last few years has enabled the deep phenotyping of cells in their native tissue context.^1–9^ Investigating the relationship between tissue structure and function using multiplexed imaging has led to important discoveries in many fields, including cancer, infectious disease, autoimmunity, and neurodegenerative disease.^10–17^ As imaging studies continue to grow in number and size, so does the need for robust computational methods for analyzing these data. In most multiplexed imaging studies, cells are the objects of interest that are quantified and investigated in downstream analyses. As such, development of methods for accurate cell annotation is an active area of research.^18–21^ Unlike assays that measure dissociated single cells such as CyTOF or single-cell RNA-sequencing, imaging data is not inherently measuring single cells and can capture substantial information content outside of cells. These extracellular features can have important biological functions. For example, the extracellular matrix is increasingly being recognized as an important modulator of the tissue microenvironment in cancer and other disease contexts.^22–25^ In addition, protein aggregates can form as extracellular deposits and have been implicated in many neurological disorders.^26^ These important acellular objects are captured in multiplexed images but are typically not the focus in multiplexed imaging studies.

One important consideration when analyzing imaging data is that the tissue sections that are captured in the images are two-dimensional cross sections of complex three-dimensional objects. Depending on the plane of tissue sectioning, what is observed in the images can be highly variable (Supplementary Fig. 1a). For example, depending on if the cell body or dendrites are in the plane of imaging, dendritic cell markers can take on a typical round cellular shape or might only be present as small spindle-like projections (Supplementary Fig. 1b). While cell segmentation methods for accurately defining the boundary of cells have recently been developed^27–29^, signal along the edges of cells can be misassigned to neighboring cells, particularly in dense tissues where cells are packed close together (Supplementary Fig. 1c-d). Furthermore, cells that are elongated or shaped distinctly from spherical cells, or anucleated cells, are difficult to capture using cell segmentation (Supplementary Fig. 1e). Identifying phenotypes at the pixel-level can address many of these issues that confound high-dimensional image analysis.

Due to these challenges with analyzing multiplexed imaging data, we developed a pipeline, Pixie, for the quantitative annotation of pixel-level features that captures phenotypes independent of traditional cell segmentation masks. We perform extensive evaluation of pre-processing steps and parameter choices to optimize clustering performance, as well as comprehensively assess stochasticity, an often-ignored aspect of high-dimensional data analysis methods. In addition, we show the application of Pixie across various tissue contexts and imaging platforms, including mass-based, fluorescence-based, and label-free technologies. Finally, we show how pixel clusters can be utilized to improve the identification of cell phenotypes. Taken together, Pixie is a complete pipeline for generating both pixel and cell-level features that is scalable, cross-platform, and publicly accessible in Jupyter notebooks that include user-friendly graphical user interfaces (GUI) for cluster adjustment and annotation.

## Results

### Overview of pixel clustering using Pixie

We created a full pipeline, Pixie, for the generation of quantitative pixel-level features from multiplexed images, with the goal of designing a pragmatic workflow that balances automation and unbiased analysis with human curation (Fig. 1). We aimed to create a robust and scalable pipeline that is user-friendly and easily extensible. After a multiplexed imaging dataset has been generated, the output is a series of images, each corresponding to a different marker of interest. The first step is to decide which markers to include in the clustering process. Typically, phenotypic markers are included, while functional markers that can be expressed across various cell types are excluded. After the user has specified the subset of markers relevant for phenotyping, we perform a series of pre-processing steps, described in detail below (Fig. 2a, Supplementary Fig. 2a). A common practice in high-dimensional data analysis is to first cluster observations into a large number of clusters (such as 100), then metacluster these clusters into biologically relevant groups.^10–12, 15, 17, 30^ This allows for the capture of rare phenotypes and more precise clusters. In Pixie, we use a self-organizing map (SOM)^31^, an unsupervised clustering algorithm, to cluster all pixels into a large number of clusters (typically 100). We then combine these into metaclusters using consensus hierarchical clustering. If necessary, the final clusters can be manually refined and annotated with biologically relevant labels using a custom-built GUI in Pixie (Supplementary Video 1). These phenotypes can be mapped back to the original images and quantified in downstream analysis. Pixie is publicly available as user-friendly Jupyter notebooks that perform all pre-processing steps and clustering, starting from multiplexed images through generation of pixel phenotype maps, where the value of each pixel corresponds to its pixel cluster.

**Figure 1:**
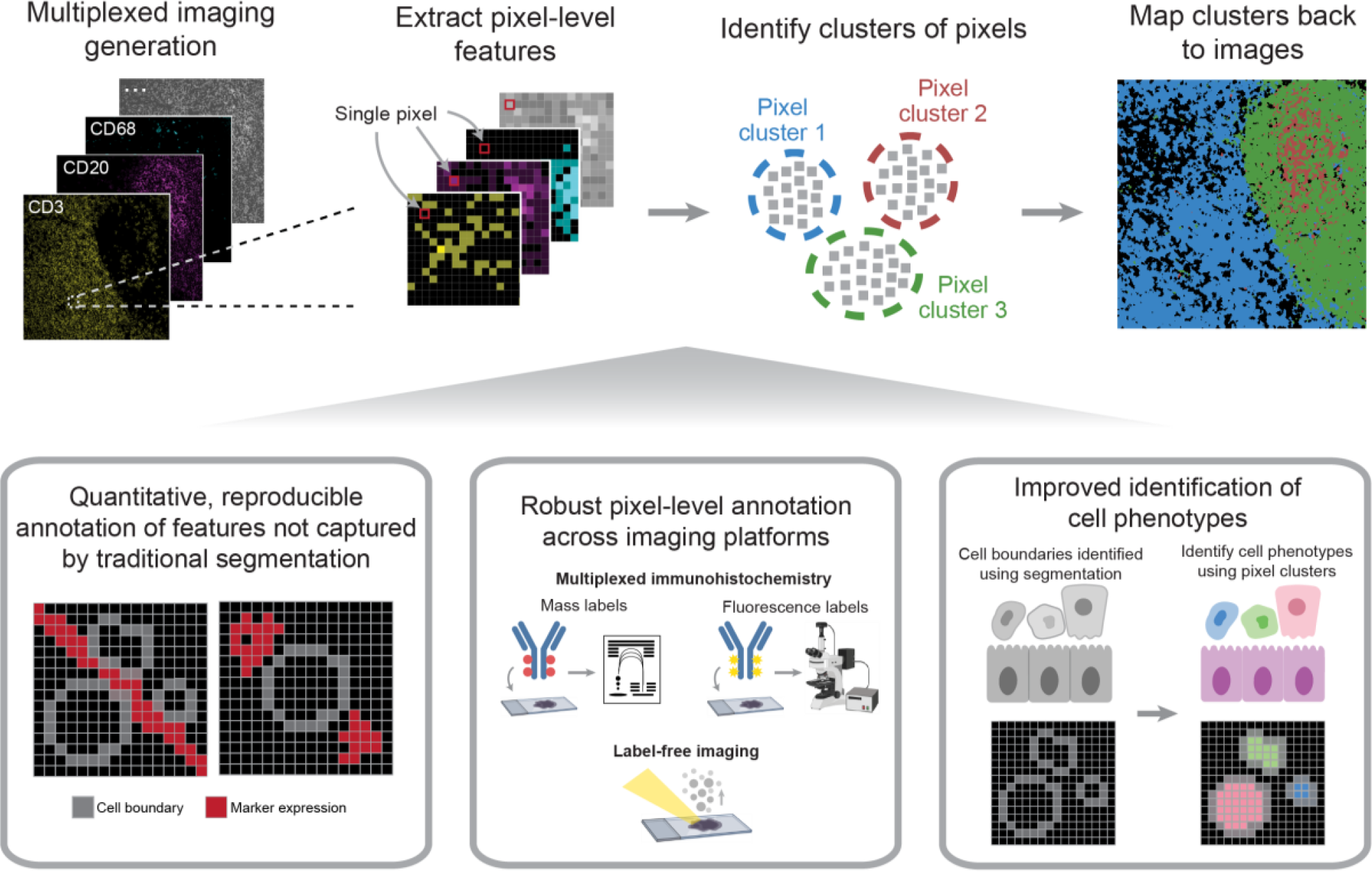
Pixie robustly captures pixel-level and cell-level phenotypes in multiplexed imaging datasets. After acquiring multiplexed images, single-pixel expression profiles are extracted from the multi-channel imaging dataset and clustered to identify pixel-level phenotypes. This method can be used to generate quantitative, reproducible annotations not captured by traditional cell segmentation, and is cross-platform and applicable across a variety of biological contexts. Finally, pixel-level phenotypes can be combined with traditional cell segmentation and be used to improve the annotation of cell-level phenotypes.

### Pixie captures the major immune phenotypes in lymph node tissue

To test pixel clustering in Pixie and optimize parameters, we used a dataset of lymph nodes stained with a panel of immune markers (Supplementary Table 1) and imaged using multiplexed ion beam imaging by time-of-flight (MIBI-TOF). In addition to T and B lymphocytes, lymph nodes contain many dendritic cells, follicular dendritic cells, and macrophages that are difficult to capture using cell segmentation (Supplementary Fig. 1b). Lymph nodes are also densely packed tissues, which confounds cell phenotyping (Supplementary Fig. 1d). Therefore, we chose this lymph node MIBI-TOF dataset for proof of principle to evaluate our method.

Using Pixie, we were able to capture the major immune phenotypes that would be expected in a lymph node (Fig. 2b). We found that automated metaclustering was able to generate accurate features and required only a small amount of manual adjustment (Supplementary Fig. 2b). Importantly, we were able to capture more fine-grained features at the pixel-level that are grouped together at the cell level. For example, CD209+CD206+ pixels clustered separately from CD163+CD206+ pixels (Fig. 2b), which were all assigned to macrophages at the cell level (Fig. 5b, Supplementary Fig 16j). Mapping these pixel clusters back to the original images, we can see that the pixel clusters accurately recapitulate underlying spatial trends in protein expression (Fig. 2c, Supplementary Fig. 3). We could clearly delineate the germinal center, B cell follicle, and surrounding T cell zone in the lymph node (Fig. 2c). Thus, we were able to quantify high dimensional phenotypes at a pixel-level using Pixie.

### Evaluating reproducibility

Many algorithms commonly used to analyze high-dimensional datasets – including unsupervised clustering methods and dimensionality reduction techniques such as tSNE (t-distributed stochastic neighbor embedding) and UMAP (uniform manifold approximation and projection) – are stochastic, meaning that there is randomness inherent to the algorithms. Similarly, a SOM is stochastic. Therefore, running the same method on the same dataset using a different random seed will generate distinct results. Good clustering results that reflect true biological phenotypes should be reproducible across different random initializations. Here, we evaluated the stochasticity of our pipeline and defined a metric, which we termed the “cluster consistency score”, for quantifying reproducibility (Methods). In an ideal situation, we would simply evaluate how consistently pixels were assigned to the same phenotype across replicates. However, like many unsupervised clustering approaches, Pixie relies on manual annotation using expert knowledge to assign each cluster to a phenotype. Given that this process would need to be repeated for each replicate in each experiment, direct evaluation of consistency in this manner would not be feasible for large numbers of tests. The cluster consistency score allows us to measure reproducibility in an automated way and quantitatively compare different parameter choices.

**Figure 2:**
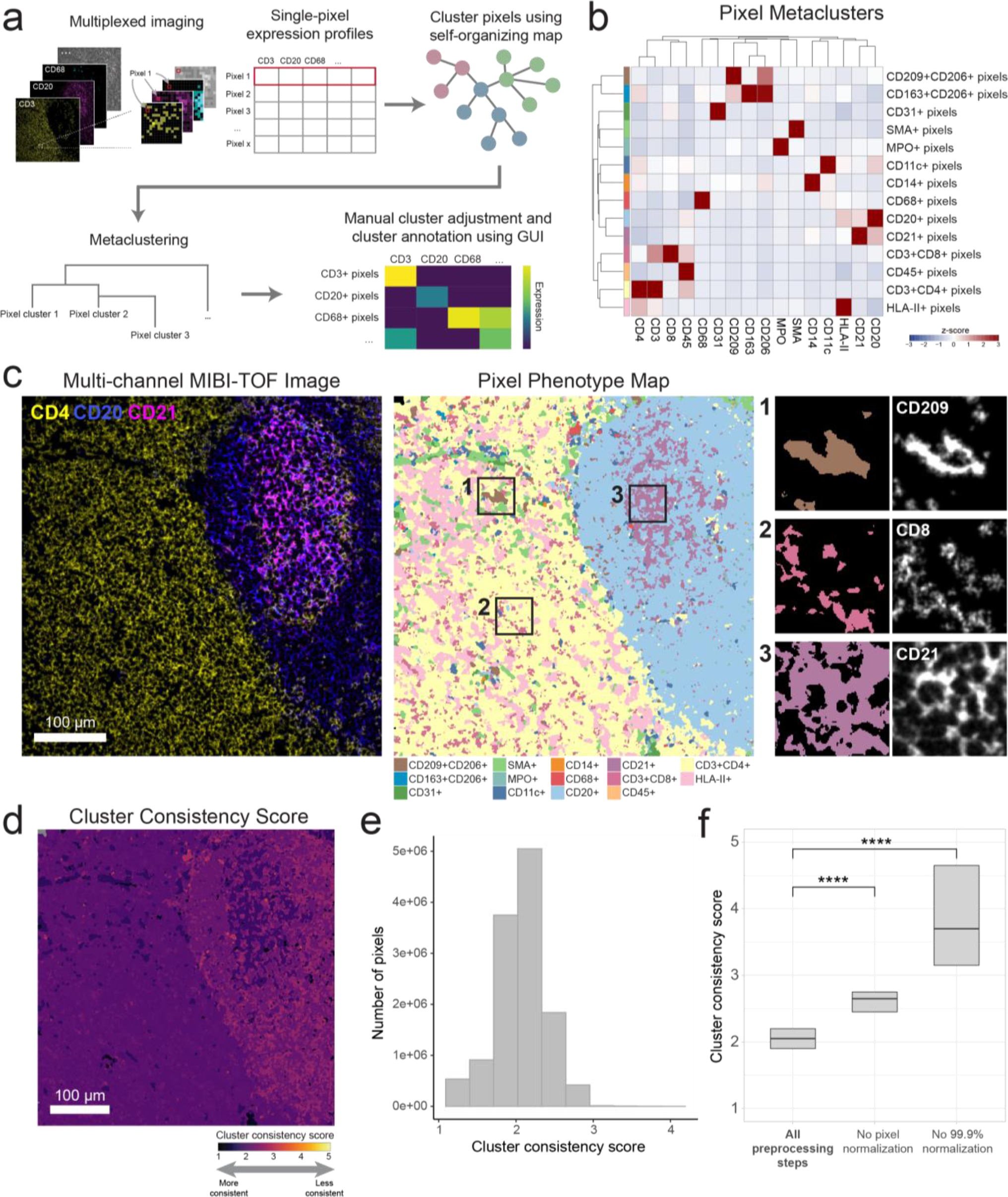
Pixie identifies accurate and consistent pixel-level features in lymph node tissue. (A) Overview of pixel clustering in Pixie. Individual pixels are clustered using a self-organizing map (SOM) based on a set of phenotypic markers. The clusters output by the SOM are metaclustered using consensus hierarchical clustering. If necessary, users can manually adjust the metaclusters, then annotate each metacluster with its phenotype based on its expression profile using our easy-to-use GUI. (B) Heatmap of mean marker expression of pixel cluster phenotypes for an example dataset of lymph node samples. Expression values were z-scored for each marker. (C) Multi-channel MIBI-TOF image of a representative field-of-view (FOV) (left), the corresponding pixel phenotype map (middle), and representative insets (right). Colors in the pixel phenotype map correspond to the heatmap in B. (D) The FOV in C colored according to the cluster consistency score. (E) Distribution of cluster consistency score across all pixels in the dataset. (F) Comparison of cluster consistency score across different pre-processing steps. **** indicates p-value < 2e-16 using a Wilcoxon test.

To calculate the cluster consistency score, we run pixel clustering on the same dataset and the same parameters five times, each time with a different random seed, then quantify how stable the cluster assignments are for each pixel across replicate runs (Methods). The cluster consistency score can be roughly interpreted as the number of different clusters a given pixel was assigned to across replicates. Lower scores indicate higher reproducibility, with a score of 1 being the best possible score. A score of 1 would indicate that in all the replicates, the same pixels were always grouped together in the same cluster. In contrast, a high score indicates bad reproducibility, meaning that the pixel was assigned to clusters that may have contained many other pixel types. We calculated this cluster consistency score for each pixel in the lymph node dataset (Fig. 2d-e), which had an overall cluster consistency score of 2.07 ± 0.32 (mean ± SD). Despite the stochastic nature of the algorithm, by viewing the pixel phenotype maps across replicate runs, we can see that the majority of pixel cluster assignments were stable across replicates (Supplementary Fig. 4a), demonstrating the reproducible nature of this method. Here, we developed a quantitative metric that can be used to assess the reproducibility of Pixie across different parameter choices.

### Optimization of Pixie for accurate pixel classification

When developing Pixie, we optimized a series of pre-processing steps that leads to more accurate, reproducible pixel clustering results (Supplementary Fig. 2a). First, we apply a Gaussian blur to the data. Akin to dropout in single-cell RNA-sequencing data where genes are not detected due to low amounts of mRNA in individual cells and inefficient mRNA capture32, multiplexed images do not capture all of the protein expressed in the tissue. Analogous to compensating for dropouts, we use a Gaussian blur to smooth the signal to make the distribution more reflective of the true underlying data. We assessed four different standard deviations for the Gaussian blur and balanced resolution of features and cluster consistency by visualizing the clusters and evaluating the cluster consistency score (Supplementary Fig. 5). We determined that a standard deviation of 2 was optimal, as it resulted in good cluster definition (Supplementary Fig. 5a-b) as well as a low overall cluster consistency score with the smallest variance (Supplementary Fig. c-e).

Next, we apply a pixel normalization step, in which for each individual pixel, we divide the signal of each marker by the total signal in that pixel, such that the sum total of that pixel is 1. The intuition behind this step is that when performing phenotyping, we are interested in the ratio between the phenotypic markers. The absolute intensity of pixels across images can be different, for example due to biological differences (such as downregulation of T cell receptors upon activation) or technical differences (drifts in instrument sensitivity, variations in tissue fixation and staining). While the underlying cause for the differences in intensity can be different, the resulting differences in absolute intensity of individual pixels confounds the phenotyping. The ratio of marker expression within each pixel contains the important phenotyping information. Without this key pre-processing step, the results contained one dominating pixel cluster that was poorly defined (low expressing for all the markers) (Supplementary Fig. 6, 7). After pixel normalization, we apply a 99.9% marker normalization step, where each marker is normalized by its 99.9th percentile value. When this step was excluded, the results contained poorly defined clusters that expressed many markers (Supplementary Fig. 8). Importantly, when either the pixel normalization or the 99.9% marker normalization steps were excluded, the cluster consistency score was significantly worse (Fig. 2f), indicating that these pre-processing steps are vital for generating consistent clusters.

Pixel-level data analysis can introduce significant computational demands. When training a SOM, all the pixels must be read into memory at the same time. For very large datasets, this is a computational bottleneck. To ensure that this approach is scalable, we wanted to see if a subsampling approach could yield equally valid results. We hypothesized that for large datasets, a random subset of pixels is a representative sample and provides the SOM with enough information to generate accurate clusters. Using a large dataset of around 800 million total pixels^33^, we trained the SOM using a random 10% subset of pixels (Supplementary Fig. 11). We found that the results of subsampling were highly concordant with the results from the whole dataset, and the cluster consistency score was consistent as well. Furthermore, we performed a similar subsampling experiment using a cyclic immunofluorescence (CyCIF) whole-slide tonsil image (27,299 pixels × 20,045 pixels).^34^ Training with a 10% subset of pixels showed highly concordant results to training with 100% of pixels (Supplementary Fig. 12). In this whole-slide image, subsampling was able to cover the data variation to generate similar clusters as training with all pixels. We have shown that for large datasets, using a random subset of pixels to train the SOM is more computationally efficient and leads to concordant results. Therefore, this approach is scalable to large datasets. Taken together, we optimized a set of pre-processing steps and parameters for consistency, biological interpretability, and scalability.

### Pixie captures extra-cellular information in multiplexed images

While cells are important features to annotate in multiplexed images, a large amount of information is captured in multiplexed images outside of cells (Fig. 3). The extracellular matrix (ECM), blood vessels, and extracellular protein aggregates are examples of objects that can exist outside of the cellular space. The biological relevance of these objects can vary based on tissue type and disease context. For example, in immune tissue such as lymph nodes (Fig. 3a), cells are densely packed within the tissue and therefore most pixels fall within cells. However, for ductal carcinoma in situ (DCIS) or triple negative breast cancer (TNBC) that contain large amounts of extracellular matrix and structural proteins (Fig. 3b-3c), pixel clustering is useful for generating a quantifiable metric for extracellular features. For these tissue types, pixels that lie outside of cells that express phenotypic markers make up a majority of the total pixels in the image (Fig. 3d). Pixie assigns a phenotype to each of these pixels, which can then be quantified and analyzed in downstream analysis.

To demonstrate how these extracellular features can be quantified, we calculated the percentage of ECM in each image of the TNBC dataset by dividing the number of pixels belonging to the ECM by the total number of pixels (Supplementary Fig. 13c-d). By comparing with MIBI-TOF images, we can see that the Pixie clusters capture different amounts of ECM in the images. We also show that there is a differential composition of ECM pixel clusters across the images. These quantifications can then be linked to various patient metadata in further downstream analysis. Importantly, pixel clustering using Pixie allows us to utilize a larger percentage of the pixels captured using multiplexed imaging technologies than cell-level analysis only.

### Pixel-level features from Pixie are reproducible across replicate MIBI-TOF runs

For multiplexed imaging approaches to be used in large translational studies and eventually in clinical diagnostics, not only must the imaging technology be robust, workflows for analyzing these data must be reproducible and accurate. In recent work by our group, we undertook a validation study to demonstrate the reproducibility of MIBI-TOF in which we assessed concordance across a dozen serial sections of a tissue microarray (TMA) of 21 cores that consisted of disease-free controls as well as multiple types of carcinomas, sarcomas, and central nervous system lesions.^35^ Here, we demonstrate the reproducibility of Pixie on replicate serial sections (Supplementary Fig. 14a).

**Figure 3:**
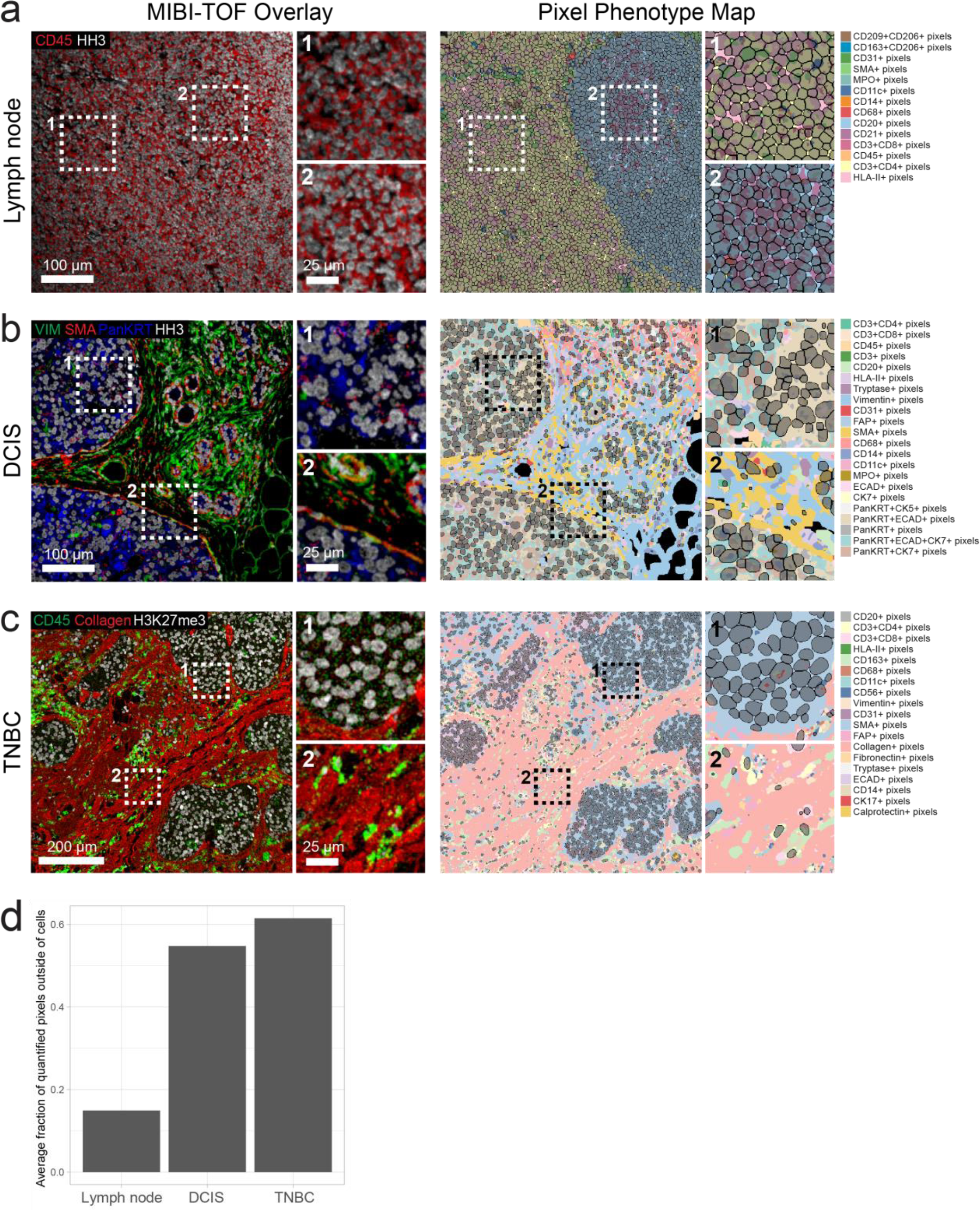
Pixie captures more information in multiplexed images than cell segmentation masks alone. Representative FOVs from (A) lymph node, (B) ductal carcinoma in situ (DCIS), and (C) triple negative breast cancer (TNBC), showing MIBI-TOF overlays (left) and pixel phenotype maps (right). Cells identified using cell segmentation are overlaid on the pixel phenotype maps in gray. Colors of the pixel phenotype maps correspond to the phenotypes indicated on the right. (D) Comparison of the average fraction of quantified pixels (pixels that were included in pixel clustering) that were outside of the cell segmentation masks across the three datasets.

Using this pipeline, we clustered the pixels across all the images into 12 phenotypes (Supplementary Fig. 14b). Because we had six serial sections per tissue core in the TMA, we could assess the reproducibility of Pixie by finding the correlation between serial sections of the same tissue core. Because true biological replicates are not possible, we compared serial sections of each tissue core as a proxy. As a result, we would expect some true biological differences between serial sections. Despite these differences, the overall Spearman correlation was high, with R2 of 0.92 ± 0.03 (mean ± SD) (Supplementary Fig. 14c), demonstrating the reproducibility of pixel clustering using Pixie. This high reproducibility was obtained despite the intensity differences across experiments, showing that our normalization pipeline is robust to technical variation and batch effects (Supplementary Fig. 14d). Here, we show that despite differences in absolute pixel intensity, Pixie is able to capture reproducible pixel phenotypes, demonstrating that pixel clustering can generate biologically meaningful annotations across entire cohorts.

### Applications of Pixie across biological contexts and imaging platforms

In one example, we used pixel-level analysis to capture biologically meaningful phenotypes in the myoepithelial layer in ductal carcinoma in situ (DCIS). In previous work, our group used MIBI-TOF to characterize the transition from DCIS to invasive breast cancer (IBC) using a 37-plex panel.^11^ DCIS is a pre-invasive lesion that is itself not life-threatening, but if left untreated, will progress to IBC in up to 50% of cases.^36^ The myoepithelial layer surrounding the ductal cells is an important histological feature that is known to undergo transformations during the progression to IBC. Normal breast myoepithelium is a thick, highly cellular layer between the stroma and ductal cells. In DCIS, the myoepithelium becomes stretched out in a thin layer with few, elongated cell bodies. In IBC, complete loss of this layer is accompanied by local invasion of tumor cells. Therefore, understanding the changes in ductal myoepithelium may shed light on what drives the progression of DCIS to IBC. Classical cell phenotyping strategies, which rely on detecting cells with a strong nuclear signal and are often optimized for conventional cell shapes, fail to capture the myoepithelial phenotype. To be able to quantify features of this important histological region, we used Pixie to identify discrete myoepithelial phenotypes (Fig. 4a). We identified seven myoepithelial phenotypes that could be quantified and compared across clinical subgroups: CK7+, CK5+, ECAD+, PanKRT+, VIM+, CD44+, SMA+. Here, we used Pixie to quantify distinct phenotypes in a small, but clinically relevant histological region that would not be captured using classical cell segmentation.

**Figure 4:**
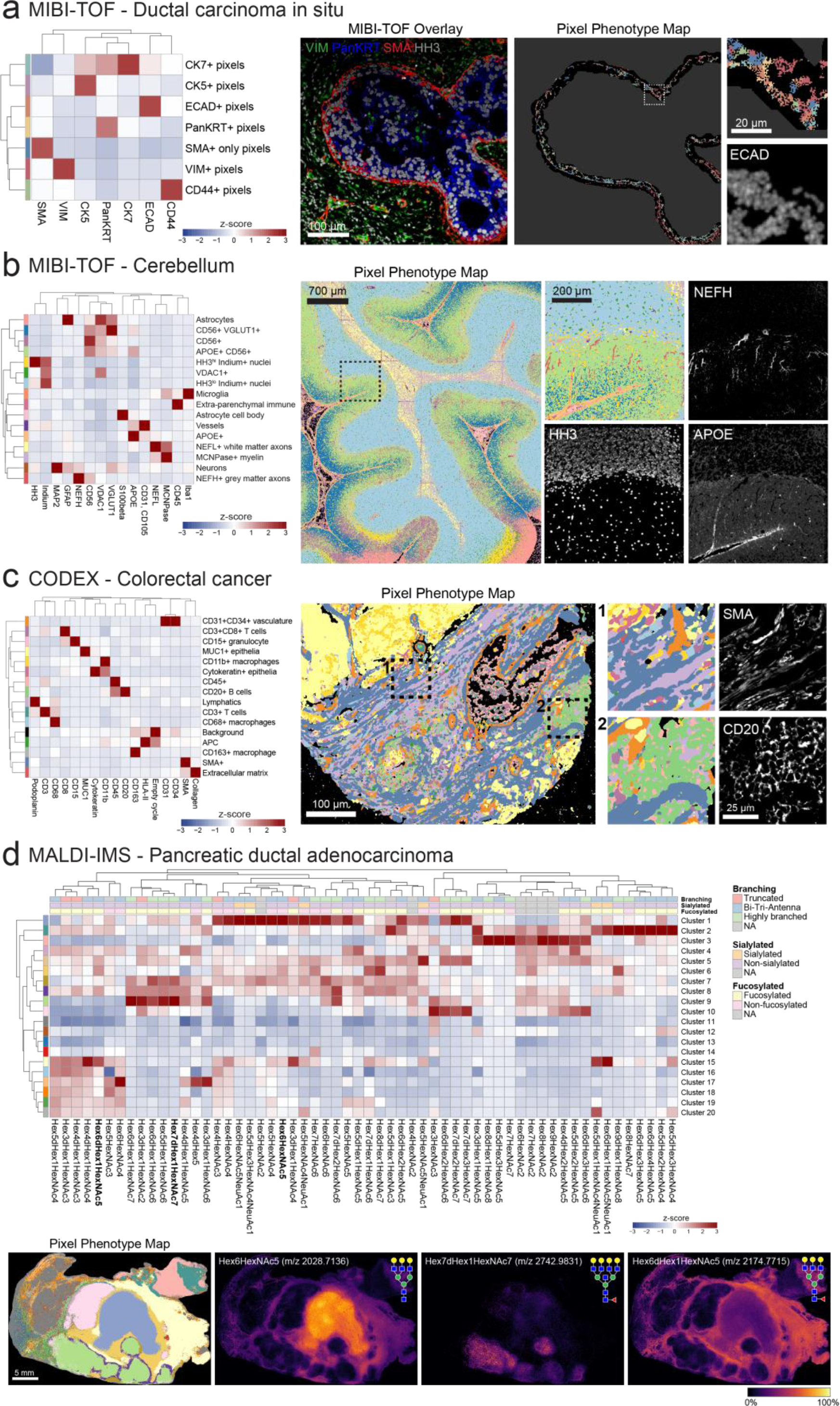
Applications of Pixie across imaging platforms and biological contexts. (A) Pixel-level phenotyping using Pixie of the myoepithelial layer in ductal carcinoma in situ (DCIS) imaged using MIBI-TOF.^11^ Heatmap of mean marker expression of the pixel clusters (left), MIBI-TOF overlay of a representative FOV (middle), and corresponding pixel phenotype map and inset (right). In the pixel phenotype map, the black region represents the myoepithelial layer. (B) Identification of pixel-level neuronal and immune features in MIBI-TOF images of the human cerebellum. Heatmap of mean marker expression of the pixel clusters (left), pixel phenotype maps of a tiled image of the cerebellum (middle), and comparison of an inset with single-marker images (right). (C) Pixel clustering in a CODEX dataset of colorectal cancer using Pixie.^17^ Heatmap of mean marker expression of the pixel clusters (left), pixel phenotype maps of a CODEX image (middle), and comparison of insets with single-marker images (right). (D) Pixel-level annotation of a MALDI-IMS dataset of pancreatic ductal adenocarcinoma using Pixie.^39^ Heatmap of mean glycan expression of the pixel clusters (top). The rows correspond to pixel clusters and columns correspond to glycans. Pixel phenotype maps (colors correspond to the heatmap) and comparison with selected glycans (bottom). Expression values were z-scored for each marker.

Another setting in which analyzing high-dimensional images has been challenging is in studies of the human brain. Studying the brain has historically been difficult due to the strong inherent autofluorescence of brain tissue, limiting the use of traditional fluorescence-based imaging techniques. We used MIBI-TOF to image neuronal and immune protein targets in the human brain. Due to the abnormal shapes of neuronal objects and the complex spatial conformations of features such as dendrites, cell bodies, and axons, classical cell segmentation techniques have limited efficacy, and detection of neuronal objects is an ongoing area of research.^13^ Using Pixie, we were able to map the full neuronal landscape of the human cerebellum, including neurons, axons, vessels, astrocytes, and microglia (Fig. 4b). For example, NEFH (neurofilament heavy chain)-expressing grey matter axons could clearly be identified as a pixel cluster. Therefore, pixel clustering using Pixie allows for the retention and classification of pixels that are independent of cell segmentation masks and can be used in downstream analysis, such as spatial analysis.

In addition, we applied Pixie to another human brain MIBI-TOF dataset, of the human hippocampus (Supplementary Fig. 15).^13^ We identified pixel cluster phenotypes corresponding to microglia, astrocytes, neurons, oligodendrocytes, vasculature, and proteopathy. To demonstrate how pixel clusters can be used in quantitative analysis, we quantified the total number of pixels of the identified Pixie clusters out of the total pixels in expert-annotated regions of human hippocampus (Supplementary Fig. 15d). We were able to observe differential pixel cluster composition across the hippocampal regions. For example, the VGLUT1^hi^CD56^lo^Tau^lo^ pixel cluster (light blue) was higher in the CA1 region, reflecting the high number of excitatory synaptic terminals in this area.^37^ The MFN2^hi^HH3^hi^ pixel cluster (purple) was highest in the dentate gyrus (DG), which may reflect a more metabolically active state of DG granular neurons.^38^ MFN2-positive neurons were also identified recently as a potential protective phenotype in the context of proteopathy development^13^, demonstrating how Pixie can be used in downstream analysis to discover biological insights.

To test the cross-platform compatibility of Pixie, we applied it to a publicly available CODEX (co-detection by indexing) multiplexed imaging dataset of colorectal cancer.^17^ In contrast to MIBI-TOF which uses metal-labelled antibodies, CODEX is a fluorescence-based method and achieves multiplexing by using DNA-barcoded antibodies and multiple cycles of imaging fluorescent nucleotides.^1^ Pixel clustering using Pixie enabled us to capture the major structural and phenotypic features in the tissue, including vasculature, epithelia, lymphatics, and immune cells (Fig. 4c). Because CODEX uses fluorescence imaging, there are significant levels of autofluorescence in the images. By including images of the empty cycles in the pixel clustering, we were able to ameliorate the effect of autofluorescence on the phenotyping by defining an autofluorescence-specific cluster. Here, we have shown the applicability of Pixie to fluorescence-based imaging approaches.

Furthermore, we applied Pixie to a previously published MALDI-IMS (matrix-assisted laser desorption ionization-imaging mass spectrometry) dataset of pancreatic ductal adenocarcinoma.^39^ MALDI-IMS is a label-free imaging approach, meaning antibodies are not used to target specific epitopes, and can be used to assess the distribution of complex carbohydrates in tissue, typified by N-linked glycans *de novo*. MALDI-IMS has been used to map N-glycan distribution across multiple cancer types.^40–42^ Unlike MIBI-TOF, CODEX, and other antibody-based approaches, MALDI-IMS was used to map glycosylation patterns and not protein expression in this dataset, so traditional image analysis techniques that rely on detecting cellular objects are not applicable. Here, we used pixel clustering in Pixie to annotate discrete phenotypes in the tissue based on glycosylation patterns (Fig. 4d). Importantly, the pixel clusters identified here were reflective of the tissue features that were manually identified in the original publication.^39^ For example, the pixel cluster localized to the center of the tissue, cluster 1 (purple-blue cluster), corresponded to the necrotic region of the tissue as determined by H&E in the original publication. One of the defining glycans in this pixel cluster was Hex6HexNAc5 (m/z 2028.7136), which was identified in the original publication as being localized to necrotic tissue. Similarly, adenocarcinoma and normal pancreatic regions were captured using pixel clustering and were defined by glycans identified in the original publication, such as Hex7dHex1HexNAc7 (m/z 2742.9831) and Hex6dHex1HexNAc5 (m/z 2174.7715), respectively. Therefore, despite the fact that MALDI-IMS is a completely different type of imaging technology that quantifies a different type of molecule on different feature scales, Pixie was able to identify histologically relevant features in an automated fashion, as well as identify glycans that were commonly co-occurring. Through these case studies, we have shown that pixel clustering can be useful across biological contexts and imaging platforms to capture pixel-level features independent of cell segmentation masks.

### Using pixel clusters to improve cell-level annotations in Pixie

Since cells are the building blocks of tissue, it is important to generate accurate annotations of cells, in addition to the pixel features described above. Development of methods for accurate cell annotations is an active area of research.^18, 19^ The current paradigm for annotating cells in images is to use unsupervised clustering, where the input features are the sum of the expression of each marker for each cell, in a manner similar to analysis of flow or mass cytometry data.^1, 10–12, 15, 17, 43^ We rely on cell segmentation to generate accurate cell masks, then integrate the expression of each marker within each cell mask to generate the expression profile for each cell. However, as discussed above, imaging data is not measuring dissociated single cells. Bright signal present along the perimeter of a cell can be inaccurately assigned to its neighboring cells, particularly in dense tissue where the cells are packed close together (Supplementary Figure 1c-d). Therefore, cell clusters using integrated expression values can have poor cluster definition due to noisy signal. In this paradigm, clusters with poor definition are usually manually inspected and compared against the images, which is a time-consuming process. Often, manual gating steps are needed to identify cells that cannot be clustered using this method. Therefore, cell phenotyping using the integrated expression of each cell requires a significant amount of manual work to visually inspect the images and adjust the clustering.

In Pixie, we use the pixel clusters resulting from the workflow described above to improve cell classification. After generating single cell masks using cell segmentation, instead of integrating the expression of each marker, we tabulate the number of pixels that belong to each pixel cluster in each cell. The number of pixel clusters in each cell is then used as the feature vector into a SOM, followed by consensus hierarchical clustering and manual cluster adjustment and annotation, as described above for pixel clustering (Fig. 5a). We hypothesized that this method would improve cell annotation because it quantifies discretized pixel phenotypes. When simply integrating the expression over the cell, misassigned pixels could have a large impact on the clustering, if their expression is very bright. Even though there may be bright pixels from a neighboring cell or from features such as dendrites that seem like they are protruding into the cell in a 2D image, the number of those pixels should be low. Because there should be few of these pixels that are misassigned to the cell of interest, the real signal should drive the clustering in this method. Furthermore, this method also quantifies the degree of protein co-expression at a pixel-level, which is information that is lost when integrating expression for each cell.

**Figure 5:**
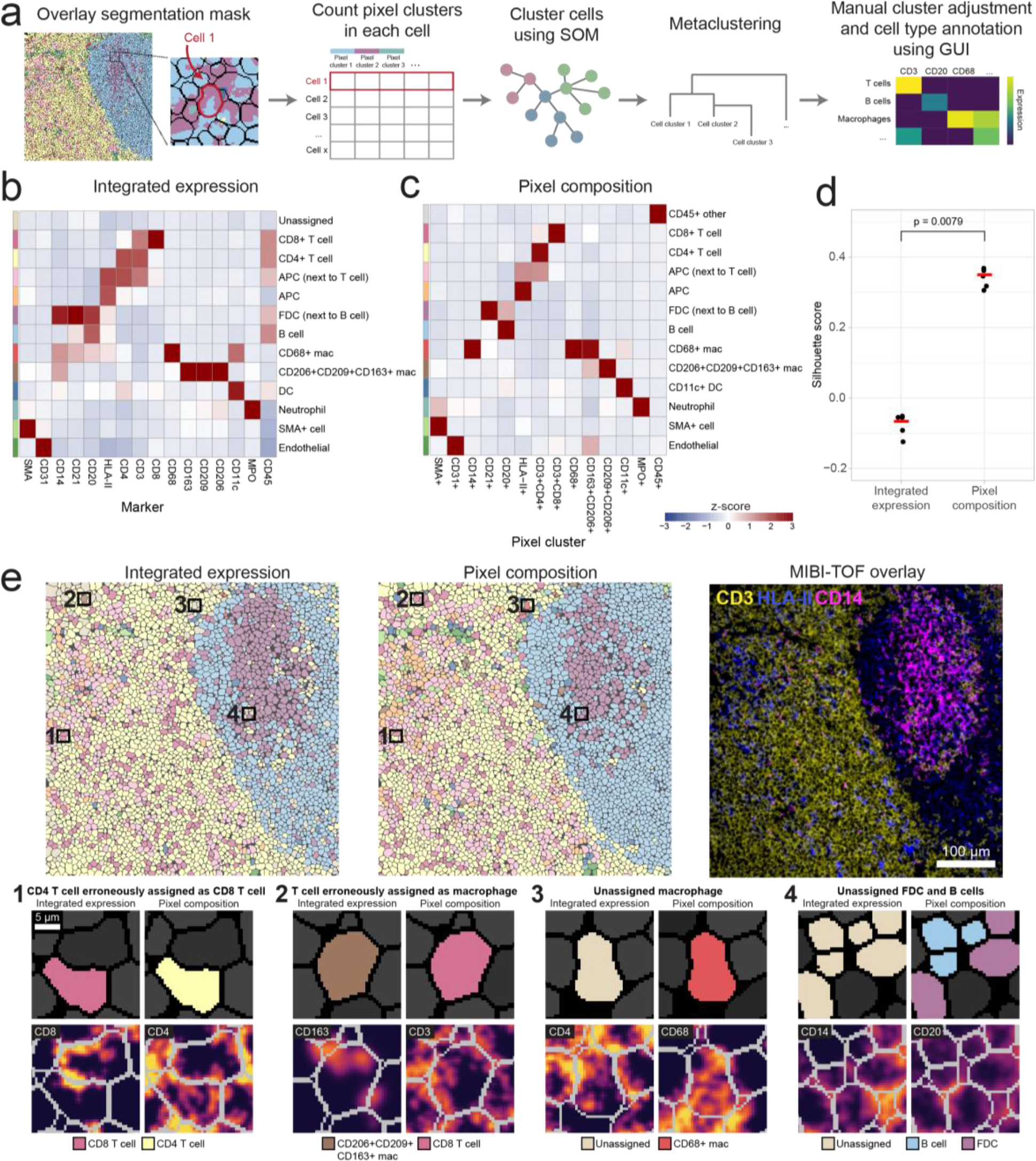
Pixel clusters can be used to improve cell annotations in Pixie. (A) Overview of cell-level phenotyping using pixel clusters. After pixel clusters have been identified and annotated, the frequency of each pixel cluster within each cell boundary (identified using cell segmentation) is used as the feature vector for cell clustering. Cells are clustered using a SOM and clusters are metaclustered using consensus hierarchical clustering. The metaclusters are then manually adjusted and annotated by the user. (B) Using the same dataset as shown in Fig. 2, heatmap of mean marker expression of cell phenotypes obtained from cell clustering using integrated marker expression for each cell. Clusters were manually adjusted and annotated. (C) Using the same dataset as displayed in Fig. 2 and in B, heatmap of mean pixel cluster frequency of cell phenotypes obtained from cell clustering using pixel cluster composition for each cell in Pixie, as outlined in A. Clusters were manually adjusted and annotated. Expression values were z-scored for each marker. (D) Comparison of Silhouette score for cell clusters obtained using integrated expression or pixel composition. Five replicates were performed for each method. Red bar indicates the average Silhouette score. P-value was determined using a Wilcoxon test. (E) Comparison of cell phenotype maps colored according to cell phenotypes obtained using either integrated expression (left) or using pixel composition (middle), and a MIBI-TOF overlay (right) for one representative FOV. Colors in the cell phenotype maps correspond to the heatmaps in B and C, respectively. Four representative examples of the advantage of using pixel composition over integrated expression for cell clustering (bottom).

To test cell clustering using Pixie, we used the lymph node dataset described above. We compared the cell expression profiles from using integrated expression and pixel composition for cell clustering (Fig. 5b-c). When using integrated expression, while the major immune phenotypes could be identified, there was one cluster that was unassigned, which was also the largest cluster (Supplementary Fig. 16c). Under this paradigm, this unassigned cluster would usually require manual comparisons with the images to determine the true phenotypes of these cells. In comparison, there was no unassigned cluster when using pixel composition to perform cell clustering. Therefore, by using pixel clusters to perform cell clustering in Pixie, we obtained cell clusters with better cluster definition and less amount of manual cluster adjustments, saving the researcher considerable time. Importantly, using pixel composition for cell clustering also resulted in significantly higher Silhouette scores, which measures how well cells are clustered with other cells that are similar to each other, a commonly used metric for evaluating clustering performance (Fig. 5d). Furthermore, the cluster consistency score was lower when using pixel composition to perform cell clustering (Supplementary Fig. 16d-e). Interestingly, after applying the same preprocessing steps (i.e. Gaussian blur, pixel normalization, 99.9% normalization) to the images before data extraction followed by clustering using integrated expression, clustering using pixel composition still resulted in a higher Silhouette score and lower cluster consistency score (Supplementary Fig. 17). While there was no unassigned cluster that was lowly expressing for all markers using the preprocessed data for integrated expression, there were many clusters that expressed ambiguous combinations of markers that would need to be further inspected (Supplementary Fig 17a).

Upon closer inspection of the images, we can identify various examples where using pixel composition to cluster cells in Pixie was advantageous to using integrated expression (Fig. 5e). In the first example, a cell was erroneously assigned as a CD8 T cell. The neighboring cell had clear CD8 expression, and the CD8 signal from this neighboring cell was confounding cell annotation. By using pixel composition, the cell was correctly assigned as a CD4 T cell. In the second example, a cell that had clear CD3 signal and was likely a T cell was being misassigned as a CD163+ macrophage, due to sparse CD163 signal that could be due to a neighboring macrophage being sectioned in 2D. Using pixel composition, this cell was assigned as a T cell. In the third example, a cell that was unassigned when using integrated expression, possibly due to noisy expression from neighboring cells, was correctly assigned as a CD68+ macrophage using pixel composition. Finally, in the fourth example, cells that were unassigned when using integrated expression were assigned as B cells and FDCs using pixel composition. Because B cells and FDCs are closely packed and often interacting in a lymph node follicle, it can be difficult to correctly assign phenotypes to these cells, emphasizing the ability of Pixie to perform well on traditionally challenging phenotypes.

While these examples demonstrate areas where Pixie is advantageous, we could also identify examples where the accuracy of the Pixie output was ambiguous (Supplementary Fig. 16h). In the first example, a cell that looked like it had clear CD3 expression (and identified as a T cell using integrated expression), also looked like it had CD20 expression, likely due to surrounding B cells, and therefore identified as a B cell using pixel composition. In the second example, while the CD20 expression was weak, the CD11c signal was strong (but more punctate), which resulted in the cells being called as dendritic cells by Pixie. Finally, there are cells in which the cells look positive for two markers that are usually not co-expressed, and the two cell clustering methods differed on the cell phenotype, such as in this example of CD4 and CD8. Taken together, while Pixie can offer advantages over the classical cell phenotyping techniques that use integrated expression, there are still examples in which the annotation is ambiguous, reflecting the difficulty of this task.

Since there is no widely accepted “gold standard” dataset where the cell phenotypes in an image are known, to more thoroughly assess the accuracy of cell clustering in Pixie, we created an expert-labeled dataset, where we manually annotated 8,068 cells across three MIBI-TOF images (Supplementary Fig. 19). Comparing with this manually annotated dataset, we showed that clustering using pixel composition in Pixie resulted in a F1 score of 0.90, while clustering using integrated expression resulted in a F1 score of 0.66. Therefore, Pixie outperformed clustering using integrated expression when comparing with human annotations.

In addition, we used Pixie to perform cell annotation of the TMA dataset described in Supplementary Fig. 14, where we assessed the concordance between serial sections of a TMA that was randomized with respect to staining and imaging day. Using the pixel clusters shown in Supplementary Fig. 14b, we classified cells into 10 cell phenotypes, then assessed concordance between serial sections by calculating the average Spearman correlation between serial sections of the same tissue core (Supplementary Fig. 21). Overall, the Spearman correlation was high, with R2 of 0.93 ± 0.05 (mean ± SD). Similar to pixel-level features, Pixie was able to capture reproducible cell phenotypes despite differences in absolute pixel intensity between imaging runs (Supplementary Fig. 21d).

Here, we have shown the utility of pixels clusters for improving cell phenotyping. By using pixel cluster composition to perform cell clustering in Pixie, we obtain accurate cell clusters that require fewer manual cluster adjustments than when using integrated expression.

## Discussion

Here, we present Pixie, a complete pipeline for identifying pixel-level and cell-level features from multiplexed imaging datasets and demonstrate its utility across a variety of tissue types and imaging platforms. Using an example dataset of lymph nodes imaged using MIBI-TOF, we demonstrate the robustness of our method. We also show applications of Pixie in DCIS and brain MIBI-TOF datasets, as well as CyCIF, CODEX, and MALDI-IMS datasets. Finally, we show how pixel clusters can be used to improve cell classification. Importantly, Pixie is available as user-friendly Jupyter notebooks that perform all steps of the pipeline and includes a GUI for manual adjustment and annotation of clusters. Our notebooks are open source can be easily customized to each user’s requirements.

Using pixel clusters to annotate features in images has been previously performed in various contexts in multiplexed imaging as well as spatial transcriptomics. In Gut et al., the authors also use a SOM to cluster pixels in iterative indirect immunofluorescence imaging (4i) images and relate these pixel features to cells, terming them Multiplexed Cell Units (MCUs).^6, 44^ In spatial transcriptomics, pixel-level features have been used to perform cell annotation and infer tissue substructures.^45–48^ Furthermore, in the spectral imaging field (e.g. mass spectrometry imaging), unsupervised pixel-level analysis has been widely used because the resolution often does not enable single-cell segmentation.^49^ Here, we show that the same preprocessing steps and pipeline that can be used to identify features in mass spectrometry imaging can also be used to identify subcellular and cellular features in other imaging technologies. Taken together, these methods demonstrate the utility of pixel-level analysis. Here, we build-upon this previous work by providing a comprehensive evaluation of the pre-processing steps and parameter choices that optimize clustering performance, show use cases across imaging platforms and biological questions, and present a user-friendly pipeline for running this method.

First, we illustrate the beneficial effect of each of the pre-processing steps in the Pixie pipeline – Gaussian blurring, pixel normalization, and 99.9% marker normalization. While we have determined an optimal parameter space for each of these pre-processing steps using multiple datasets, these parameters may need to be tuned for each individual dataset. For example, we determined a lower Gaussian blur was appropriate when analyzing the smaller myoepithelial area in DCIS and that no Gaussian blur was needed when analyzing MALDI-IMS data. There are many other parameters of the SOM that could be tuned, such as the learning rate, initialization of the cluster centers, and distance function. We found that the default parameters of the SOM (see Methods) worked well for all of our example use cases captured using different technologies (i.e. MIBI-TOF, CODEX, CyCIF, MALDI-IMS) and at different resolutions. However, if necessary, these could be easily changed in our pipeline. We encourage users to visualize the resulting pixel clusters alongside marker expression images to assess the best parameter choices for their own datasets. Furthermore, while we have shown that a subsampling approach for dataset sizes up to 800 million total pixels is able to generate highly concordant clusters as training with the full dataset, as the size of imaging datasets continue to grow, further subsampling experiments may be necessary.

It is important to note that Pixie is intended for images that have been pre-processed to remove as many imaging artifacts as possible. Since many imaging artifacts can be technology-specific, many imaging technologies have developed platform-specific image preprocessing pipelines.^1, 50–52^ Pixie is not intended for background subtraction or noise removal, and is instead intended for the detection of acellular and cellular objects from images with a high signal-to-noise ratio. Furthermore, because of the pixel normalization step, Pixie cannot detect phenotypes characterized by differential expression of the same marker. To be able to detect phenotypes with differential expression, one possibility is to do a second clustering step after Pixie. For example, after identifying a CD4+ pixel cluster using Pixie, users can do an additional clustering step on just the pixels within this pixel cluster to identify a high and low population.

In Pixie, the user specifies the number of metaclusters for consensus hierarchical clustering and can manually adjust these metaclusters (Supplementary Fig. 2b). While there have been various methods developed to computationally determine what the algorithm predicts is the optimal number of metaclusters, we found that manual inspection of the cluster expression profiles is a fast and accurate method for determining the number of relevant phenotypes. The number of relevant phenotypes, although subjective, may vary based on the biological question of interest and is best understood by the researchers leading the study. For example, for a study that is focused on granular subsets of myeloid cells, it may be important to stratify populations that are expressing combinations of CD206, CD209, CD163, CD68, CD14, and CD16 within the monocyte/macrophage lineage. However, for another study that is interested in mapping the general immune landscape, these markers may be grouped together into the macrophage population. Analogous to “human-in-the-loop” approaches in the fields of machine learning and artificial intelligence, the manual annotation step allows us to utilize the biological expertise of the user to improve the results more quickly. Furthermore, one of the reasons that it has previously been difficult to manually adjust clustering results is that there was no good way to manually interact with the clustering outputs. Importantly, manual adjustments and annotations can be easily made using a custom-built GUI in our Jupyter notebooks (Supplementary Video 1).

While we used a SOM as the clustering algorithm in our pipeline, there are many other unsupervised clustering algorithms that have been developed for a similar purpose.^53–56^ We chose to use a SOM because it is accurate, fast, and scalable, which is particularly important for this method because we are clustering a large number of pixels in which the number of observations can approach 1 billion. In contrast, the number of cells from single-cell RNA-sequencing or CyTOF experiments are usually on the order of thousands or millions. One popular clustering algorithm is the Leiden algorithm, which is built upon the Louvain algorithm, both often used in transcriptomic analysis and implemented in the popular Seurat package.^57, 58^ PhenoGraph, another popular clustering algorithm, is a graph-based method that identifies communities using Louvain. We performed a time comparison of a SOM (implemented in FlowSOM), Leiden (implemented in Seurat), and PhenoGraph (implemented in Rphenograph) and observed that a SOM has the fastest runtime (Supplementary Fig. 22). If the user wishes to use another clustering algorithm, the modular nature of our code allows the clustering algorithm to be easily replaced.

Additionally, we have shown that Pixie can quantify biologically meaningful features that are not captured by traditional cell segmentation across disease contexts and imaging platforms. In one example, we used Pixie to define a clinically meaningful feature in DCIS that could stratify patient groups. One of the defining features of DCIS is that the myoepithelium becomes stretched out as the tumor cells proliferate and expand. We found that normal breast myoepithelium exists in a luminal, E-cadherin (ECAD)-positive phenotypic state, which transitions to a more mesenchymal, vimentin-positive state in DCIS, which aligns with an analogous shift in tumor cell differentiation. In previous work from our group, we built a random forest classifier using 433 parameters for predicting which DCIS patients would progress to IBC, including the pixel-level features.^11^ Importantly, a high abundance of the ECAD+ myoepithelium pixel cluster was the number one predictor of IBC recurrence in this study, highlighting the utility of this pixel clustering method for discovering biological insights.

Lastly, we demonstrate the utility of using pixel clusters to annotate cell-level features and show its improvement over using integrated marker expression for cell annotation. While difficult to quantify in an automated manner, in our group’s experience, using pixel clusters to perform cell clustering requires fewer manual cluster refinement steps. While we offer one improvement to the traditional cell clustering methodology, there are many other algorithms that have been developed to address similar problems. REDSEA (Reinforcement Dynamic Spillover EliminAtion) improves cell assignments by correcting for spillover signal at cell boundaries.^20^ While the pixel clustering method described here similarly accounts for spillover signal from neighboring cells, Pixie also accounts for pixels that may not be at the cell boundary that are confounding accurate classification, such as noisy pixels or pixels from objects actually not associated with the cell of interest, such as dendrites from nearby cells or the extracellular matrix. Furthermore, while Pixie relies on unsupervised clustering, another class of algorithms that can perform cell type assignment relies on feeding the algorithm prior knowledge.^18, 19, 21, 46, 59^ Astir is a probabilistic model that uses prior knowledge of marker proteins to assign cells to cell types in multiplexed imaging datasets.^18^ Recently published algorithms, CELESTA and STELLAR, identify cell types in multiplexed images by utilizing spatial information.^19, 21^ One of the inputs of CELESTA is a user-defined cell type signature matrix and STELLAR relies on annotated reference datasets. Methods that rely on a priori knowledge limit the potential for discovery of cell states and relies on an accurate reference list of marker expression. Although we currently do not make use of the spatial location of pixels or cells for performing phenotype annotation, this represents an exciting new avenue for future work.

As the amount of multiplexed imaging datasets continues to grow, automated, fast, and scalable approaches for analyzing these data are needed. Pixie is a simple, fast method that can generate quantitative annotations of features both independently and in conjunction with cell segmentation that will enable the comprehensive profiling of various tissues across health and disease.

## Methods

### Pixel clustering methodology

The pixel clustering method in Pixie is illustrated in Supplementary Fig. 2a and described above. Single pixel expression profiles were extracted from single-channel TIFs – all pixels from all fields-of-view (FOVs) in a dataset are run in aggregate. Pixels that had zero expression for all clustering markers were excluded. The data was then Gaussian blurred with a standard deviation of 2 (unless otherwise noted), pixel normalized by dividing by the total pixel expression, and 99.9% marker normalized by dividing by each marker’s 99.9^th^ percentile. The 99.9% normalization step is necessary because markers that are systematically brighter would otherwise likely drive the clustering. By normalizing all markers to their 99.9^th^ percentile, markers have a more even contribution to the clustering results. Next, we used a SOM to cluster all pixels into 100 clusters. Unless otherwise noted in the manuscript, we used the following parameters of the SOM for all clustering: grid size = 10 × 10, start learning rate = 0.05, end learning rate = 0.01, initialization function = random, distance function = Euclidean, training passes = 10. Next, the mean expression profile of each of the 100 clusters was determined, z-scored for each marker, then z-scores were capped to a maximum value of 3. The clusters were then metaclustered using consensus hierarchical clustering using the z-scored expression values. Metaclusters were manually adjusted and annotated using a custom-built GUI (Supplementary Video 1). These final phenotypes were mapped back to the original images to generate pixel phenotype maps. To generate expression heatmaps, we calculated the mean expression for each cluster and found the z-score for each marker.

All processing was performed on a Google Cloud Compute Engine instance. The machine type, number of cores, and available memory were adjusted based on the size of the dataset.

There are many parameters in the SOM that can be tuned. As described above, Pixie uses a SOM to first overcluster the pixels, then metaclusters. For the lymph node dataset, we tested using a SOM to directly cluster pixels into 15 clusters (Supplementary Fig. 9). This resulted in poor pixel cluster definition for some clusters, as well as a worse cluster consistency score, indicating that initial overclustering is a critical step for accurate results. Furthermore, the number of passes through the dataset that is used to train the SOM is another tunable parameter. For this dataset, we compared training the SOM using 1 pass, 10 passes, and 100 passes (Supplementary Fig. 10). We found that 10 passes were sufficient to achieve accurate clusters (Fig. 2b-c), and that increasing the number of passes up to 100 did not change the clustering results.

We also compared our unsupervised clustering method to image thresholding using Otsu’s method. For the lymph node dataset, we used Otsu’s method to classify each pixel as positive or negative for each of the markers. We show that counting the number of positive markers in each pixel leads to a very large number of possible phenotypes. For example, if using 16 markers, pixels that are defined by one positive marker have 16 possible phenotypes (i.e. each of the markers). However, pixels that are defined by three positive markers (e.g. CD14+CD11c+CD20+) have 473 total phenotypes, while pixels that are defined by six positive markers (e.g. CD14+CD4+CD3+CD68+CD8+CD206+) have 991 total phenotypes (Supplementary Fig. 20c). It is not trivial to assign each pixel to a small set of phenotypes using thresholding. In contrast, the output of Pixie is a small, defined set of phenotypes, where each pixel is assigned to one of these phenotypes.

### Cell clustering methodology

Cell segmentation for all datasets was performed using the pre-trained Mesmer segmentation model.^27^ We used histone H3 as the nuclear marker, and a combination of CD45, CD20, and HLA-II as the membrane marker. For each cell in the image that was identified using Mesmer, we counted the number of each pixel cluster in each cell. We normalized these values by the total cell size and applied a 99.9% feature normalization (features here were the pixel clusters). Cells were then clustered using a SOM, and metaclustered using consensus hierarchical clustering, analogously to pixel clustering as described above.

In addition, we also performed segmentation using a combination of ilastik and CellProfiler as described previously (Supplementary Fig. 18).^51^ We show that using these segmentation masks, there is still a large low-expressing unassigned cluster using integrated expression for clustering. We also show that clustering using pixel composition had a higher Silhouette score.

### Cluster consistency score

To assess the stochasticity of Pixie, we created the “cluster consistency score” metric. Across different replicate runs of the same input data, the same phenotype may be output as a different cluster number, so assessing reproducibility by comparing the number of pixels belonging to the same phenotype is not easily automated and instead requires significant amounts of manual annotation. For example, the pixel cluster that is defined by CD20 may be pixel cluster 1 in the first run and pixel cluster 2 in the second run. Manual annotation of each cluster is infeasible for large numbers of tests when assessing pre-processing steps and different parameter choices. To measure reproducibility quickly and quantitatively, we created the cluster consistency score. Cluster consistency score is calculated as follows:

1. For one set of parameters, we run the entire pipeline using the same input data five times, each time with a different random seed. We call these replicates 1-5.
2. For each replicate, for each cluster, we quantify the minimum number of clusters in another run that it takes to get to 95% of the pixels in that cluster. For example, if there are 1000 pixels belonging to cluster 1 of replicate 1, for these pixels, we count the number of pixels in each cluster of replicate 2. We then rank this count table and determine the minimum number of clusters in replicate 2 it takes to get to 950 pixels in cluster 1 of replicate.
3. For a single replicate, we calculate this number in a pairwise manner with all other replicates. For example, for replicate 1, we calculate this number for replicate 1-replicate 2, replicate 1-replicate 3, replicate 1-replcate 4, replicate 1-replicate 5. These numbers are averaged.
4. Steps 2 and 3 are repeated for each cluster in each replicate. The result is that each pixel cluster in each replicate has a score associated with it.
5. These scores are mapped back to the pixel assignments. For example, if a pixel was assigned as pixel cluster 1 in replicate 1, the corresponding score determined in the previous step is assigned to that pixel. Each pixel is assigned 5 features, corresponding to the score from each replicate. These 5 features are averaged for each pixel, resulting in one score for each pixel.

See code for full implementation. A low score indicates good reproducibility while a high score indicates bad reproducibility, meaning that the pixel was assigned to clusters that may have contained many other pixel types. At the cell level, the same paradigm was used, but instead of pixel clusters, we assessed cell clusters.

We selected 95% when calculating the cluster consistency score because we determined that it was a good benchmark value. As expected, lowering this threshold resulted in lower cluster consistency scores, and raising this threshold resulted in higher cluster consistency scores (Supplementary Fig. 4b).

To demonstrate that the cluster consistency score calculated using five replicates is a stable measurement, we performed 100 tests of this calculation, where each test consisted of five replicates (Supplementary Fig. 4f). Across the 100 tests, we found that the mean cluster consistency score was 2.18 ± 0.12, showing that the mean cluster consistency score was stable across the 100 tests.

While it is computationally prohibitive to compute the cluster consistency score using a large number of replicates, to show an example of the reliability of the cluster consistency score, we ran the Gaussian blur standard deviation comparison using 100 replicates (Supplementary Fig. 5e). As explained above, we perform pairwise comparisons, so 100 replicates equates to 4,950 comparisons. Compared to the calculation done using five replicates, the conclusions are the same when performing the calculation with 100 replicates. The cluster consistency score was highest for no Gaussian blur, while a Gaussian blur of 2 had a low cluster consistency score as well as the smallest variance.

### Benchmarking cluster consistency score with reference cell datasets

To benchmark the cluster consistency score using high-dimensional datasets that are commonly analyzed using stochastic methods, we used two publicly available single cell datasets, a CyTOF dataset of whole blood and single-cell RNA-sequencing dataset of peripheral blood mononuclear cells (PBMCs) (Supplementary Fig. 4c-e).^60, 61^ Since imaging data is not inherently single cell, there can be spatial overlap between neighboring cells (Supplementary Fig. 1c-d) and can be confounded by segmentation inaccuracies. Pixels can also be thought of as sparse samples of a whole cell, because it is only a fraction of the total cell volume. Therefore, we would expect better reproducibility for single-cell datasets generated using dissociated single cells than for image-based features. Even for these single cell datasets clustered using FlowSOM and Leiden respectively, the cluster consistency score was above 1, showing that stochasticity is an inherent feature of various unsupervised clustering algorithms used in multi-dimensional data analysis that should be taken into account.

The CyTOF dataset contained 1,140,035 cells from whole blood and was downloaded from: https://doi.org/10.5281/zenodo.3951613. We randomly subsampled 5000 cells from the dataset and clustered the cells into 100 clusters using FlowSOM and metaclustered into 15 metaclusters using consensus hierarchical clustering. The single-cell RNA-sequencing dataset was downloaded from the Seurat tutorial website. The Seurat dataset contained 2,700 peripheral blood mononuclear cells (PBMCs) and was downloaded from: https://cf.10xgenomics.com/samples/cell/pbmc3k/pbmc3k_filtered_gene_bc_matrices.tar.gz. The data was processed as outlined here: https://satijalab.org/seurat/articles/pbmc3k_tutorial.html. The data was log normalized and the first 10 PCs from PCA were used as the input features. We constructed a KNN graph based on the Euclidean distance in PCA space and used the Leiden algorithm to cluster cells. For both CyTOF and RNA-seq datasets, the cluster consistency score was calculated as outlined above.

### Lymph node MIBI-TOF dataset

Lymph nodes from six patients were imaged using a MIBI-TOF instrument with a Hyperion ion source using 37 markers (Supplementary Table 1). 12 fields-of view (FOVs) were imaged at a field size of 500 μm × 500 μm at 1,024 × 1,024 pixels.

### Replicate serial section TMA dataset

The dataset assessing the reproducibility of TMA serial sections was previously published in Liu and Bosse et al.^35^ The dataset included 21 different tissue cores, including various disease-free tissue, carcinomas, sarcomas, and central nervous system lesions. 165 FOVs were imaged at a field size of 500 μm × 500 μm at 1,024 × 1,024 pixels. The processed imaging data is available at https://doi.org/10.5281/zenodo.5945388. Imaging parameters and pre-processing methodology (background subtraction, denoising) are described in the manuscript. Pixel clustering and cell clustering were performed as described above. Markers included in the clustering are indicated in Supplementary Fig. 14b.

### Decidua MIBI-TOF dataset

The decidua MIBI-TOF dataset was previously described in Greenbaum, Averbukh, Soon et al.^33^ The dataset contained 222 FOVs imaged at a field size of 800 μm × 800 μm at 2,048 × 2,048 pixels. Imaging parameters and pre-processing methodology (background subtraction, denoising) are described in the manuscript. To compare the full dataset against a subset dataset, we randomly subsampled 10% of the total number of pixels for each replicate run. Subsequent steps (Gaussian blur, pixel normalization, 99.9% marker normalization) were performed as described above. Markers included in the clustering are indicated in Supplementary Fig. 11a-b. **Tonsil CyCIF dataset** The whole-slide tonsil CyCIF dataset was previously described in Schapiro et al.^34^ The whole slide image was 27,299 × 20,045 pixels. The data was downloaded at https://www.synapse.org/#!Synapse:syn24849819/wiki/608441. To compare the full dataset against a subset dataset, we randomly subsampled 10% of the total number of pixels for each replicate run. Subsequent steps (Gaussian blur, pixel normalization, 99.9% marker normalization) were performed as described above. Markers included in the clustering are indicated in Supplementary Fig. 12a-b.

### DCIS MIBI-TOF dataset

The DCIS MIBI-TOF dataset was previously published in Risom et al.^11^ The dataset contained 168 FOVs imaged at a field size of 500 μm × 500 μm at 1,024 × 1,024 pixels. The processed imaging data is available at https://data.mendeley.com/datasets/d87vg86zd8/3. Imaging parameters and pre-processing methodology (background subtraction, denoising) are described in the manuscript. Processing steps (Gaussian blur, pixel normalization, 99.9% normalization) were performed as described above. Markers included in the clustering are indicated in Supplementary Fig. 13a.

For the myoepithelial analysis in Fig. 4a, masks of just the myoepithelial zone were generated as described in the manuscript. Images were first subset for pixels within the myoepithelial masks, then pixels within the myoepithelium mask were further subset for pixels with SMA expression > 0. Upon inspecting clustering results for a few different standard deviations (sigma) for the Gaussian blur, we determined that a Gaussian blur of 1.5 was more appropriate for this use case, since we were interested in discrete pixel features in a small histological region of the full image. Subsequent steps (pixel normalization, 99.9% marker normalization) were performed as described above.

### TNBC MIBI-TOF dataset

This dataset will be published in a forthcoming publication, currently under preparation. TNBC samples were imaged using a MIBI-TOF instrument. The dataset used in this manuscript contained 31 FOVs imaged at a field size of 800 μm × 800 μm at 2,048 × 2,048 pixels. Processing steps (Gaussian blur, pixel normalization, 99.9% normalization) were performed as described above. Markers included in the clustering are indicated in Supplementary Fig. 13b.

To generate the manually labeled dataset for benchmarking cell clustering, we selected three representative FOVs from the TNBC dataset. For each cell, we manually called each cell as positive or negative for each marker using the QuPath software. A total of 8,068 cells were labeled for 21 markers. From these, we manually assigned each cell to a cell phenotype. We used scikit-learn in Python to calculate the F1 scores.

### Cerebellum MIBI-TOF dataset

This dataset will be published in a forthcoming publication, currently under preparation. Human cerebellum samples were imaged using a MIBI-TOF instrument. Each FOV was 700 μm × 700 μm at 1,024 × 1,024 pixels, and 42 FOVs were tiled to generate the final cerebellum image. Processing steps (Gaussian blur, pixel normalization, 99.9% normalization) were performed as described above. Markers included in the clustering are indicated in Fig. 4b.

### Hippocampus MIBI-TOF dataset

The hippocampus MIBI-TOF dataset was previously published in Vijayaragavan and Cannon et al.^13^ 196 FOVs imaged at a field size of 400 μm × 400 μm were tiled together to form an image of 11,264 pixels × 8,704 pixels. The processed imaging data is available at https://doi.org/10.25740/tx581jb1992. Imaging parameters and pre-processing methodology (background subtraction, denoising) are described in the manuscript. Processing steps (Gaussian blur, pixel normalization, 99.9% normalization) were performed as described above. Markers included in the clustering are indicated in Supplementary Fig. 15b.

### Colorectal cancer CODEX dataset

The CODEX dataset was previously published in Schürch et al.^17^ The processed imaging data was obtained from The Cancer Imaging Archive at https://doi.org/10.7937/tcia.2020.fqn0-0326. We selected 20 representative FOVs from CRC_TMA_A: reg012_X01_Y01_Z09, reg039_X01_Y01_Z08, reg059_X01_Y01_Z11, reg046_X01_Y01_Z09, reg015_X01_Y01_Z08, reg052_X01_Y01_Z09, reg047_X01_Y01_Z08, reg027_X01_Y01_Z09, reg035_X01_Y01_Z09, reg018_X01_Y01_Z09, reg042_X01_Y01_Z08, reg041_X01_Y01_Z09, reg069_X01_Y01_Z09, reg063_X01_Y01_Z08, reg068_X01_Y01_Z09, reg024_X01_Y01_Z09, reg019_X01_Y01_Z09, reg064_X01_Y01_Z10, reg061_X01_Y01_Z10, reg045_X01_Y01_Z10. Images were 1,440 pixels × 1,920 pixels. Processing steps (Gaussian blur, pixel normalization, 99.9% normalization) were performed as described above. In addition to the markers indicated in Fig. 4c, empty cycle TIFs were also included in the clustering.

### Pancreatic ductal adenocarcinoma MALDI-IMS dataset

The MALDI-IMS dataset was previously published in McDowell et al.^39^ Raw MALDI-IMS data (corresponding to Figure 3 in the original publication) was provided upon request by Dr. Richard Drake. Data was provided as mis, bak and tsf files, which were imported into SCiLs Lab 2022a imaging software. In SCiLs Lab, N-glycan spectra were normalized by total ion count and converted to vendor-neutral imzML format.^62^ The imzML and ibd files were parsed using pyimzML in Python, and the expression at each m/z peak was extracted as single-channel TIF images corresponding to each extracted m/z peak. The images were 670 pixels × 438 pixels. These m/z peaks were then mapped to glycans by accurate mass as annotated in the original paper. These single-channel TIFs were then processed and clustered as described above for single-marker MIBI-TOF images. Upon inspecting clustering results for a few different standard deviations (sigma) for the Gaussian blur, we determined that no Gaussian blur was necessary for pixel clustering of MALDI-IMS data. This is expected because MALDI-IMS data is lower resolution than MIBI-TOF data. Subsequent steps were performed as described above.

### Visualization

Plots were created using the ggplot2 and pheatmap R packages and the matplotlib Python package. Schematic representations were created with biorender (https://biorender.io/). Figures were prepared in Adobe Photoshop and Adobe Illustrator.

## Data availability

User-friendly Jupyter notebooks for running Pixie are available at https://github.com/angelolab/pixie. The code used to generate the figures is available at https://github.com/angelolab/publications/tree/main/2022-Liu_etal_Pixie

## Acknowledgements

We thank Dr. Richard Drake for providing the MALDI-IMS data, Dr. Zaza M. Ndhlovu for providing the lymph node tissue, and Meelad Amouzgar for feedback on the manuscript. C.C.L was supported by the National Institute of Allergy and Infectious Diseases of the NIH under award number F31AI165180 and the Stanford Graduate Fellowship. N.F.G. was supported by NCI CA246880, NCI CA264307, and the Stanford Graduate Fellowship. E.F.M. was supported by the National Science Foundation (graduate research fellowship grant 2017242837) and training grant 5T32AI007290. K.X.L. was supported by the A∗STAR National Science Scholarship. M.A. was supported by the National Institutes of Health (grants 5U54CA20997105, 5DP5OD01982205, 1R01CA24063801A1, 5R01AG06827902, 5UH3CA24663303, 5R01CA22952904, 1U24CA22430901, 5R01AG05791504 and 5R01AG05628705), the Department of Defense (contracts W81XWH2110143), the Wellcome Trust and other funding from the Bill and Melinda Gates Foundation, Cancer Research Institute, the Parker Center for Cancer Immunotherapy and the Breast Cancer Research Foundation.

## Ethics declaration

M.A. is an inventor on patents related to MIBI technology. M.A. is a consultant, board member, and shareholder in Ionpath Inc.

## Author Contributions

C.C.L., N.F.G., and M.A. developed the methodology. C.C.L., A.K., and S.R.V. developed the software. E.F.M. and D.M. stained and acquired the lymph node and cerebellum MIBI-TOF data, respectively. K.X.L. assisted in analyzing the MALDI-IMS data, D.M. assisted in analyzing the cerebellum data, and B.J.C. assisted in analyzing the hippocampus data. J.L.R. created the manually annotated cell phenotype dataset. C.C.L. wrote the manuscript. All authors reviewed the manuscript and provided feedback. M.A. supervised the work.

