## SupplementaryInfo for "Robust phenotyping of highly multiplexed tissue imaging data using pixel-level clustering"

**Supplementary Figure 1: Challenges with analyzing multiplexed imaging data**

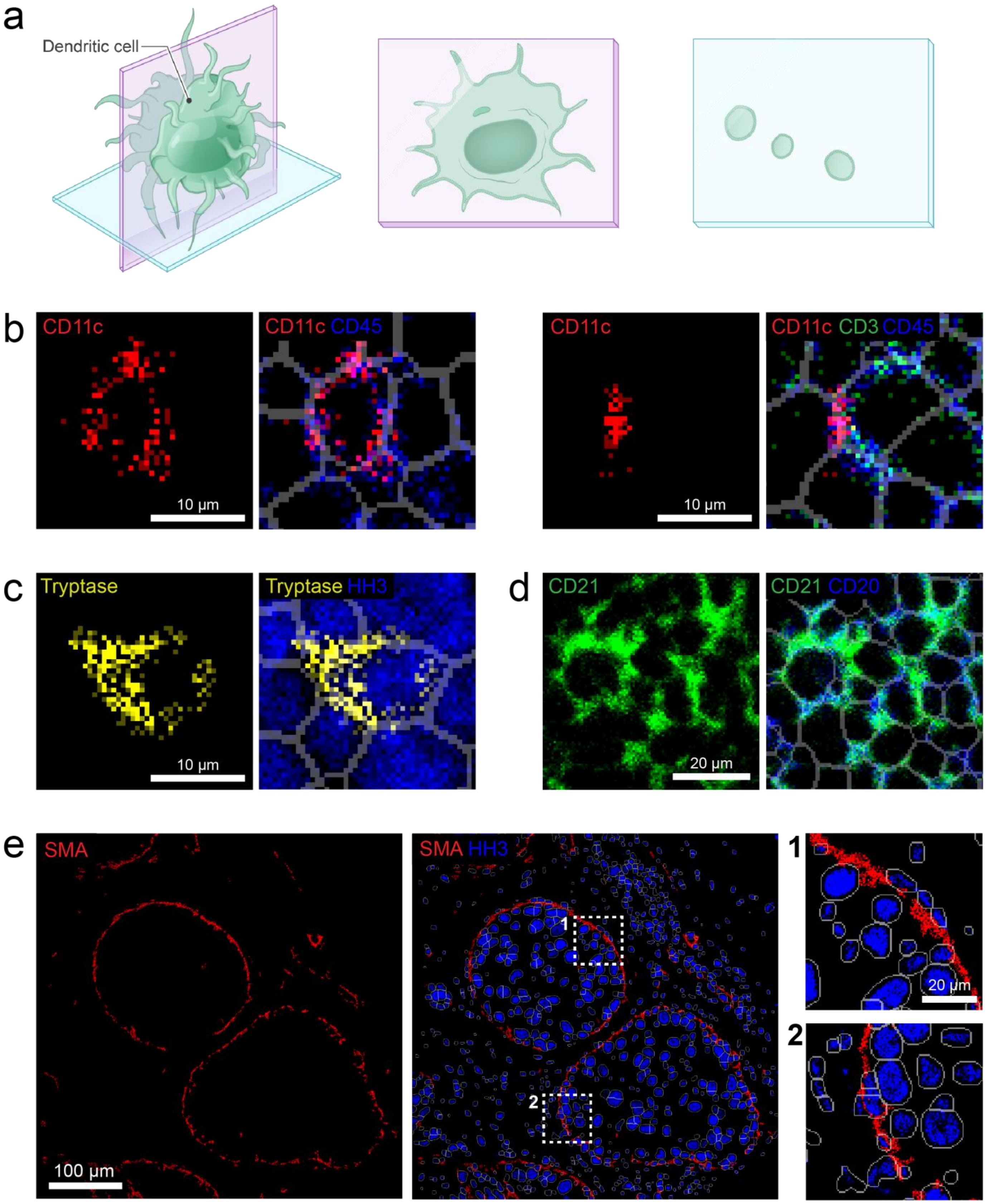


(A) Illustration of sectioning a 3D object in 2D. The objects that are viewed on the slide are highly dependent on the sectioning plane. (B) Example of a dendritic cell marker, CD11c, appearing cellular (left) or as an acellular object (right).

Gray lines correspond to cell boundaries obtained using segmentation. (C) Example of a marker overlapping into the neighboring cell. A mast cell marker (tryptase) is shown here. (D) Example of two overlapping markers when tissue is dense, and cells are in close contact with each other. A follicular dendritic cell and B cell marker (CD21) and B cell marker (CD20) are shown here. (E) Example of a feature not captured by traditional cell segmentation. The thin myoepithelial layer surrounding the ductal cells in ductal carcinoma in situ (DCIS) is shown here.

**Supplementary Figure 2: Pixel clustering pipeline in Pixie**

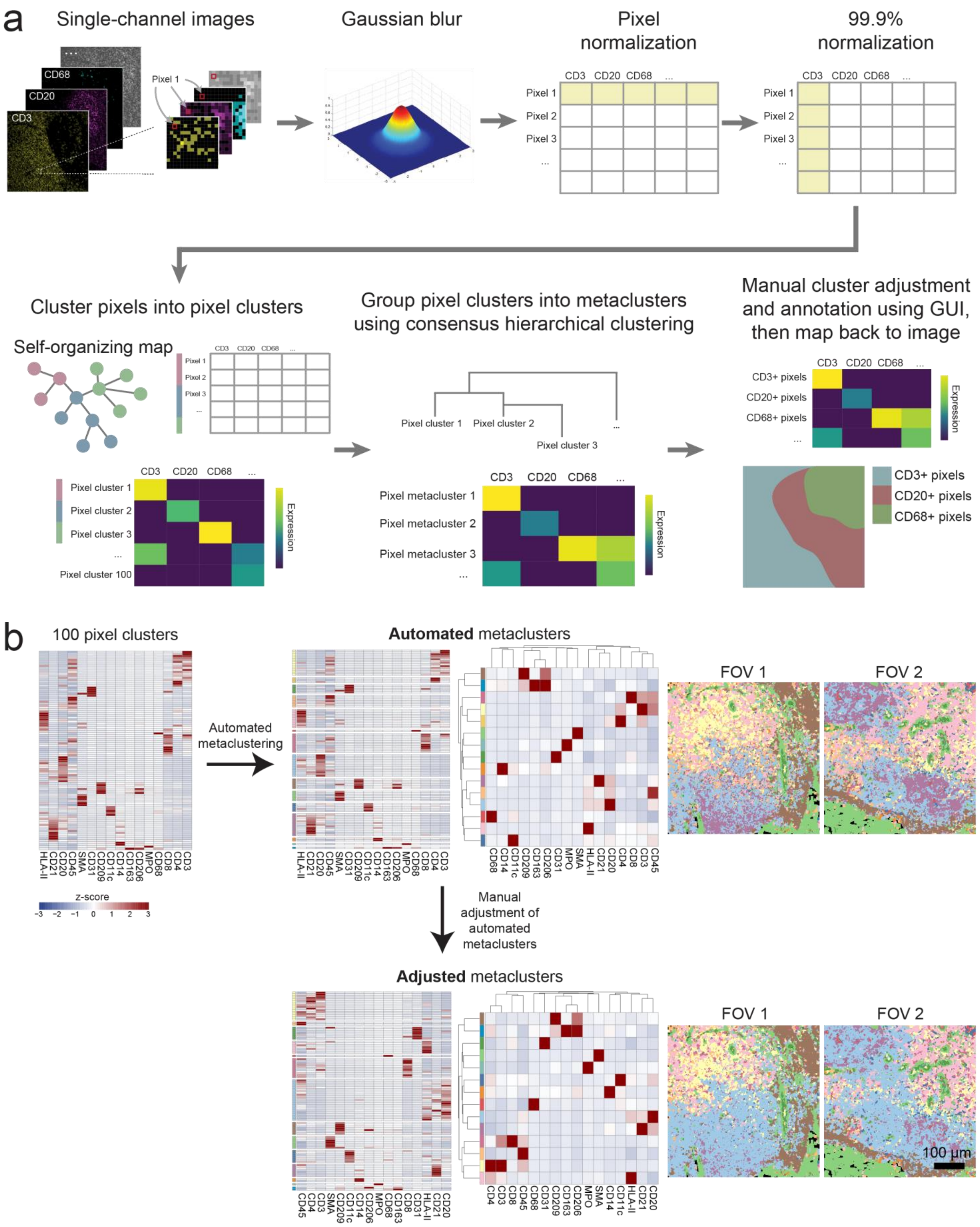

(A) Detailed overview of the pixel clustering pipeline in Pixie. First, pixel expression profiles are extracted from the images. A Gaussian blur, pixel-level normalization, and 99.9% marker normalization are applied. The transformed pixels are then clustered using a self-organizing map (SOM). The clusters output by the SOM are metaclustered using consensus hierarchical clustering. Finally, the user can manually adjust the metaclusters and annotate each metacluster with its phenotype based on its expression profile using our easy-to-use GUI. These final pixel clusters can then be mapped back to the original images. (B) Comparison of manual metacluster adjustment. After SOM clustering, the 100 clusters are metaclustered using consensus hierarchical clustering, which is a fully automated process (top). The assignment of clusters to the metaclusters can then be manually adjusted to better reflect expected biology (bottom). Pixel phenotype maps for automated metaclusters and adjusted metaclusters are shown (right).

**Supplementary Figure 3: Additional examples of pixel clustering**

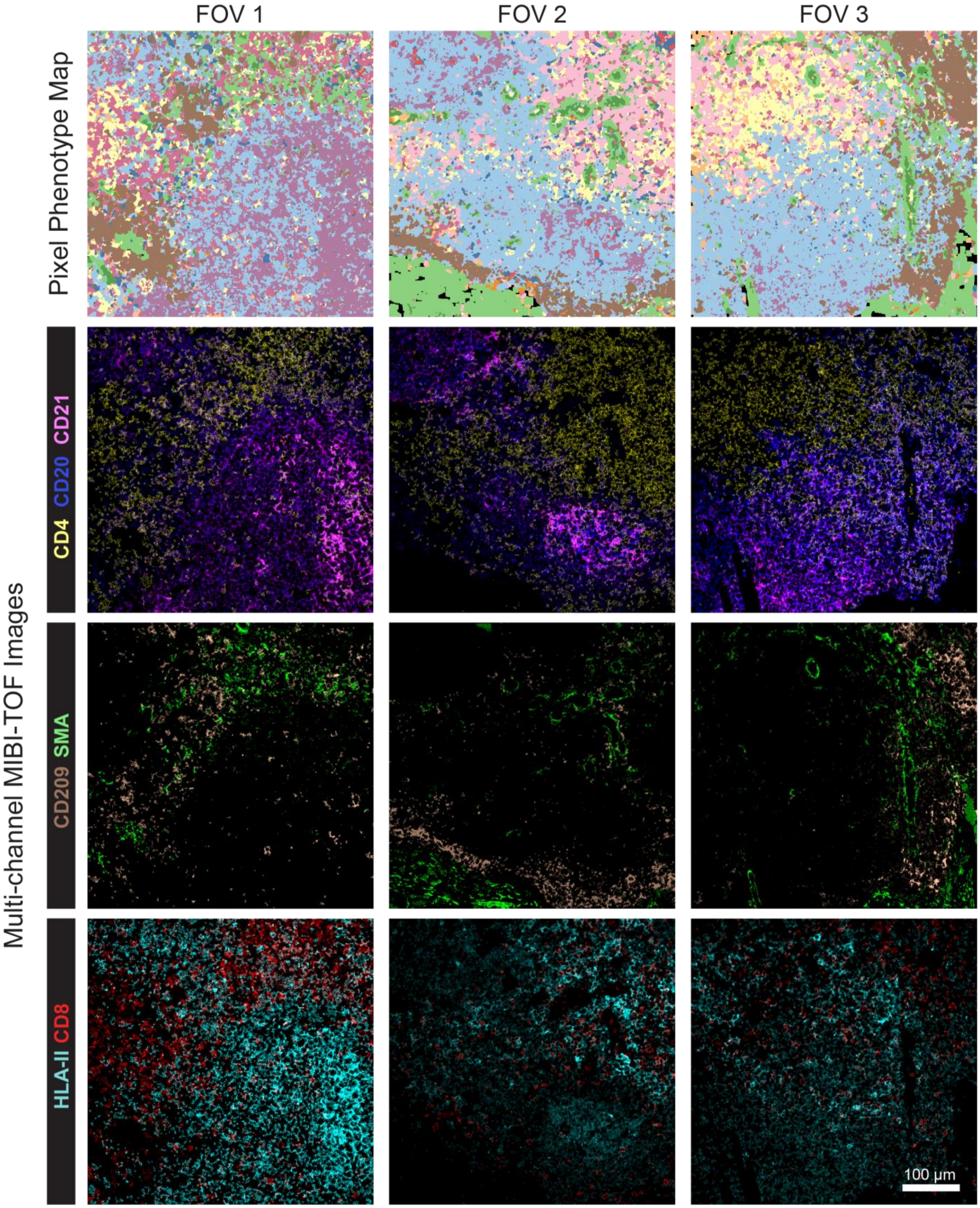


Additional examples of pixel clustering of the lymph node dataset. Top row shows the pixel phenotype maps, where the colors correspond to the heatmap in Fig. 2b. The second, third, and fourth rows show MIBI-TOF overlays for various markers. Each column is one FOV.

**Supplementary Figure 4: Assessing reproducibility**

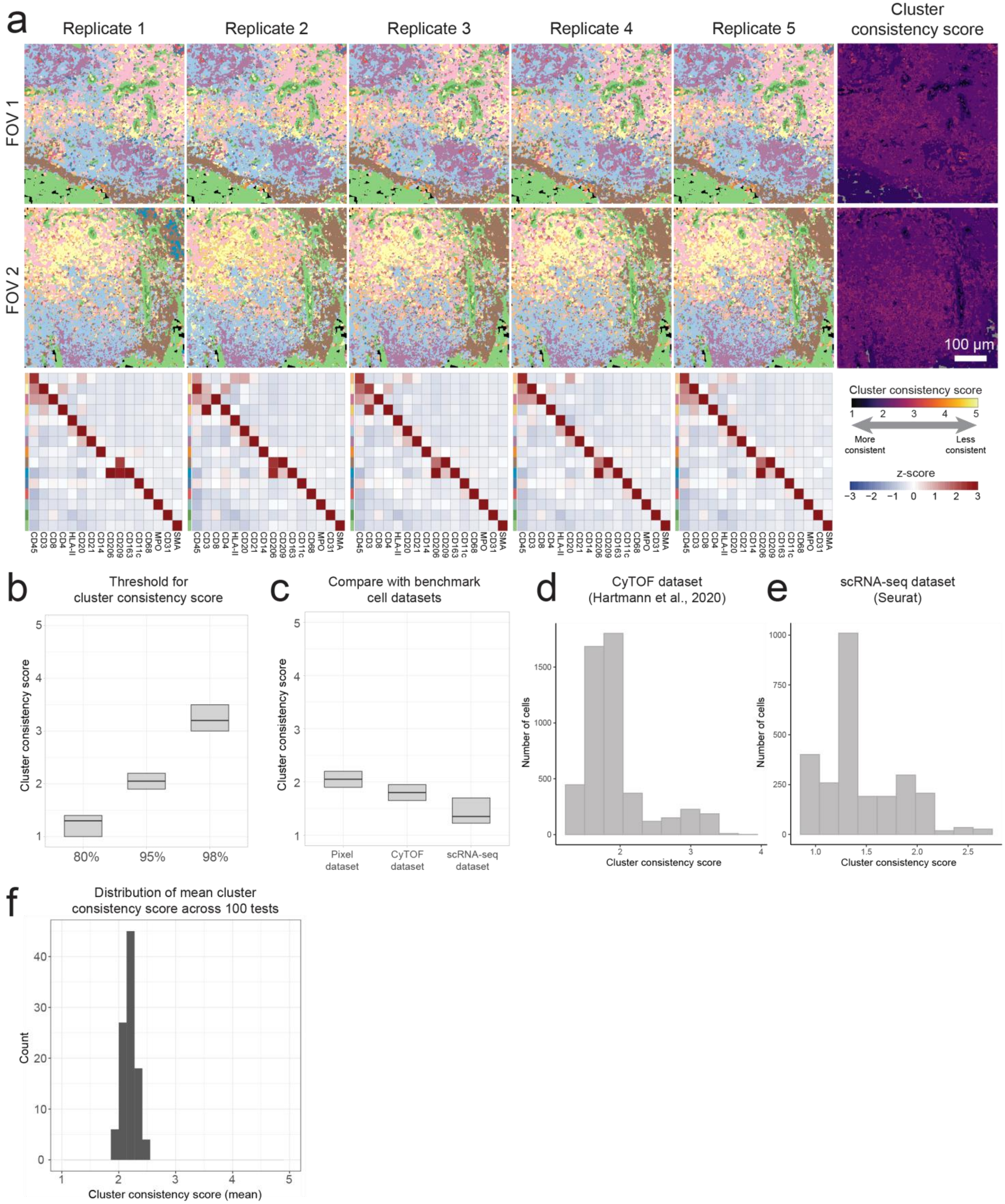


(A) Pixel phenotype maps of five replicates of the same FOV (each replicate initialized with a different seed). Pixel phenotype maps colored according to the heatmaps in the bottom row. The right column shows the same FOV colored according to cluster consistency score. (B) Distribution of cluster consistency score across all pixels for different thresholds for calculating the cluster consistency score. (C) Comparison of distribution of cluster consistency scores of pixel clustering of the lymph node dataset (all pre-processing steps) and two benchmark cell datasets, a reference CyTOF dataset and single-cell RNA-seq dataset. (D) Distribution of cluster consistency score for a reference CyTOF dataset.[60] (E) Distribution of cluster consistency score for a reference single-cell RNA-sequencing dataset (2,700 PBMC dataset from the Seurat tutorial website).[61] (F) Distribution of mean cluster consistency score across 100 tests, where each test comprised of 5 replicates of the SOM trained on the same dataset.

**Supplementary Figure 5: Gaussian blur**

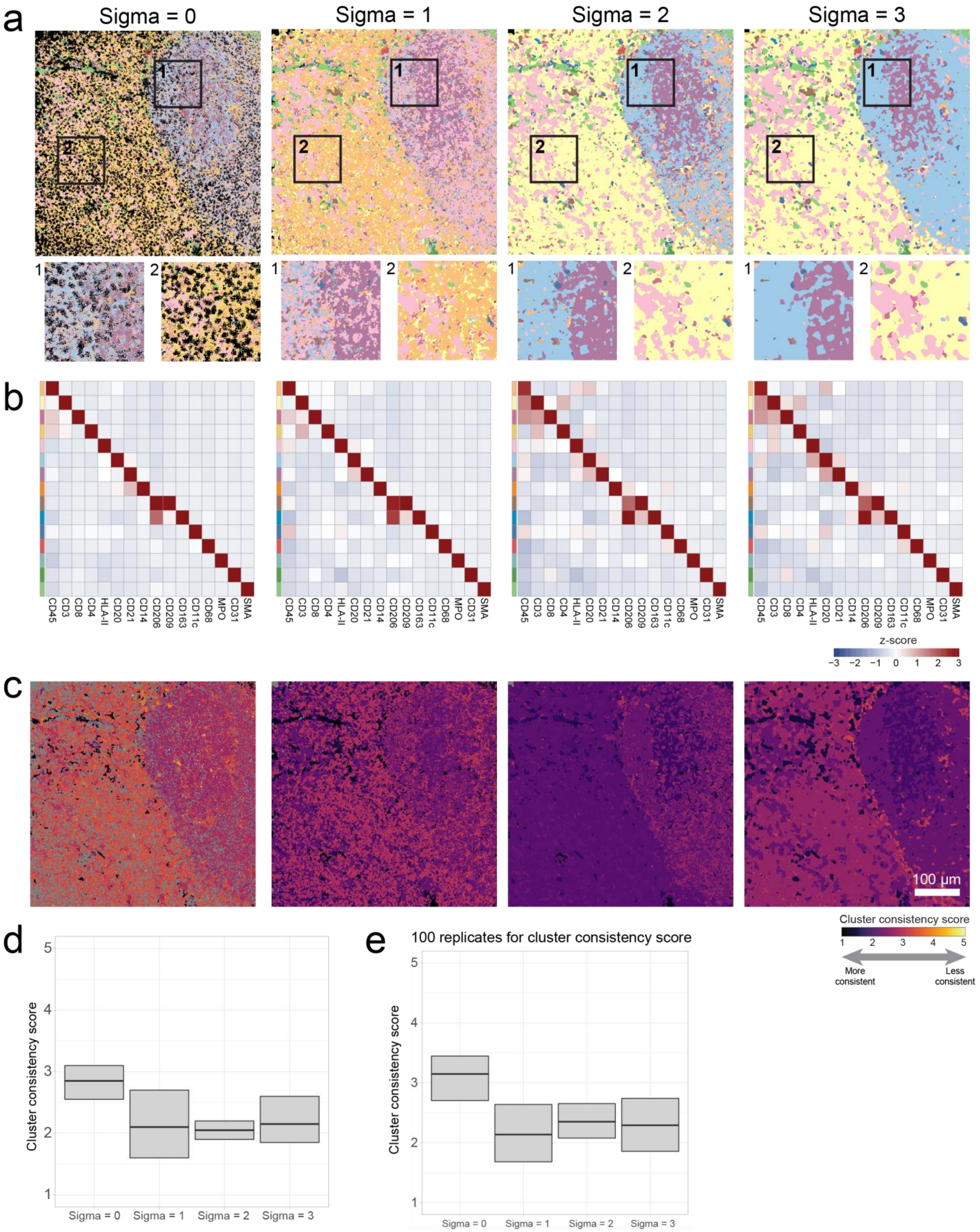

(A) Pixel phenotype maps showing the effect of increasing the standard deviation (sigma) for the Gaussian kernel of the blur. (B) Heatmaps of the pixel cluster expression profiles corresponding to each sigma. Colors of the color bars correspond to the pixel phenotype maps in A. (C) FOVs colored according to cluster consistency score for each sigma. (D) Comparison of the distribution of cluster consistency scores across all pixels in the dataset for each sigma. (E) Comparison of the distribution of cluster consistency scores across all pixels where 100 replicates were used for the cluster consistency score calculation.

**Supplementary Figure 6: No pixel normalization**

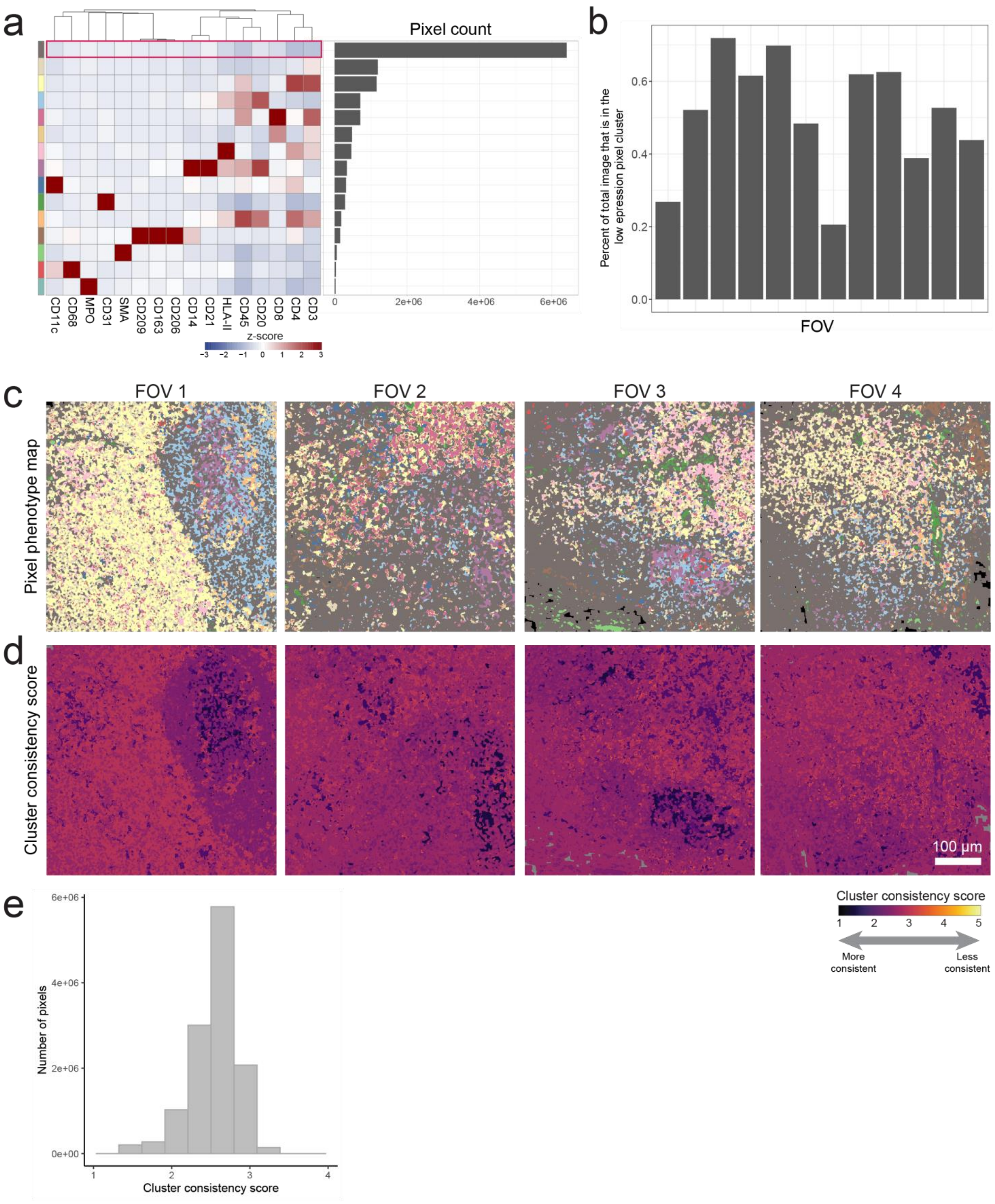


(A) Heatmap of mean marker expression of pixel cluster phenotypes where there was no pixel normalization performed. The number of pixels per cluster is shown on the right. (B) The percentage of the total pixels in each image that were assigned to the low expression pixel cluster. (C) Pixel phenotype maps for four representative FOVs (colored according to the heatmap in A). (D) The same FOVs in C colored according to cluster consistency score. (E) The distribution of cluster consistency score across all pixels in the dataset.

### Supplementary Figure 7: No pixel normalization in additional datasets

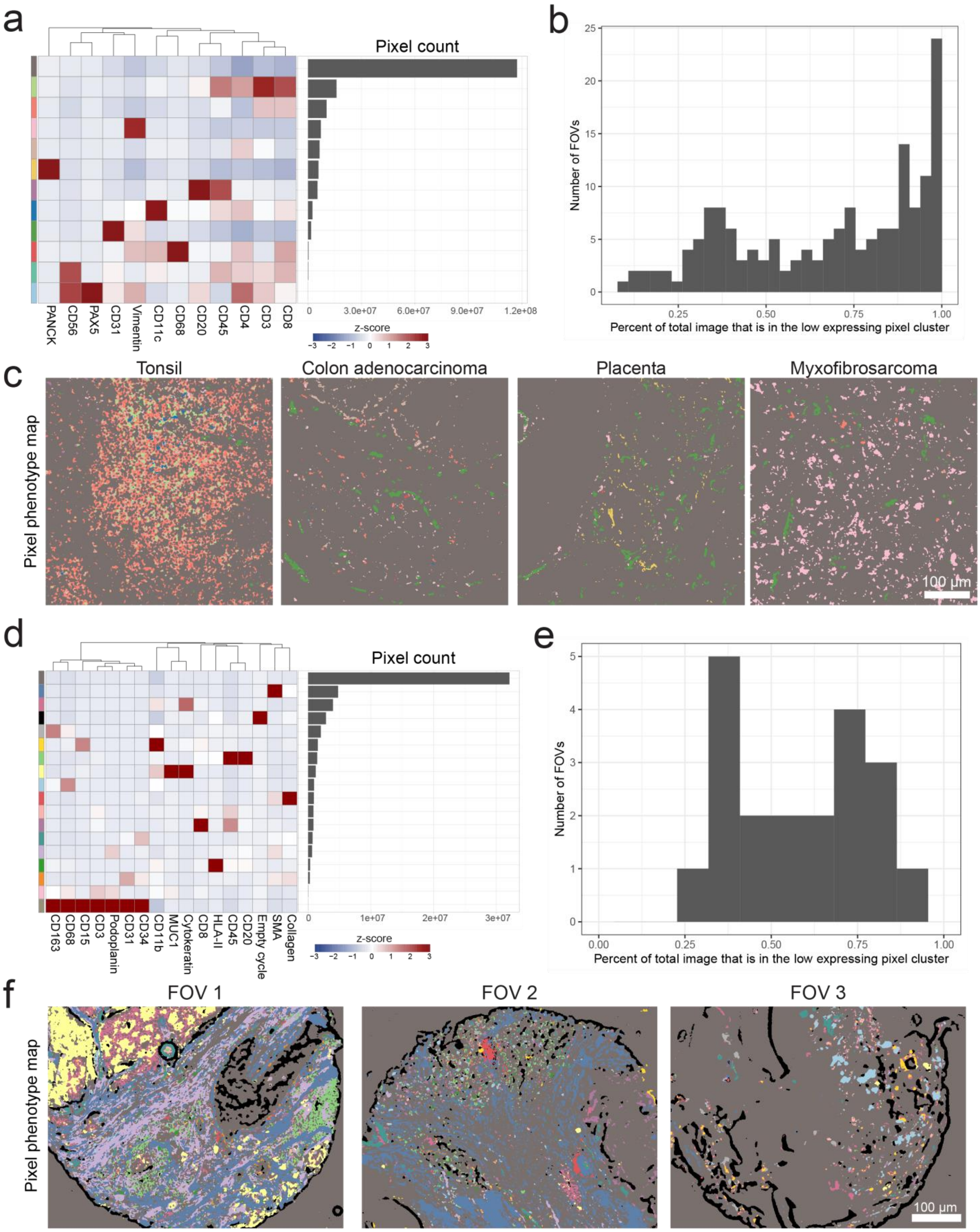

(A-C) Pixel clustering with no pixel normalization applied to MIBI-TOF dataset of 21 different tissue cores.[35] (D-F) Pixel clustering with no pixel normalization applied to CODEX dataset of colorectal cancer.[17] (A, D) Heatmap of mean marker expression of pixel cluster phenotypes where there was no pixel normalization performed. The number of pixels per cluster is shown on the right. (B, E) The distribution of the percentage of the total pixels in each image that were assigned to the low expression pixel cluster. (C, F) Pixel phenotype maps for representative FOVs (colored according to the heatmap in A or D, respectively).

**Supplementary Figure 8: No 99.9% normalization**

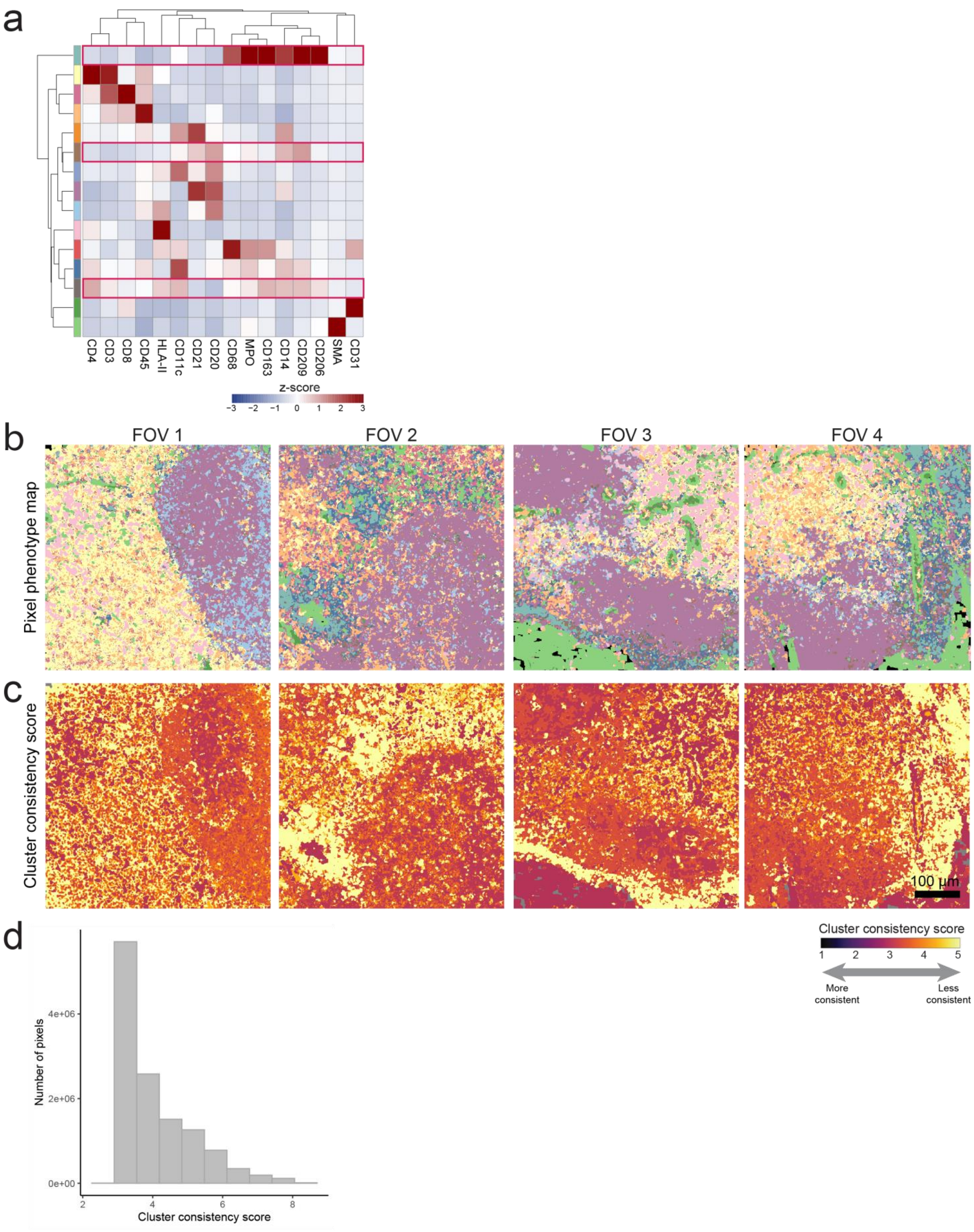

(A) Heatmap of mean marker expression of pixel cluster phenotypes where there was no 99.9% marker normalization performed. The red boxes indicate ambiguous pixel clusters with poor cluster definition. (B) Pixel phenotype maps for four representative FOVs (colored according to the heatmap in A). (C) The same FOVs in B colored according to cluster consistency score. (D) The distribution of cluster consistency score across all pixels in the dataset.

**Supplementary Figure 9: SOM with 15 nodes**

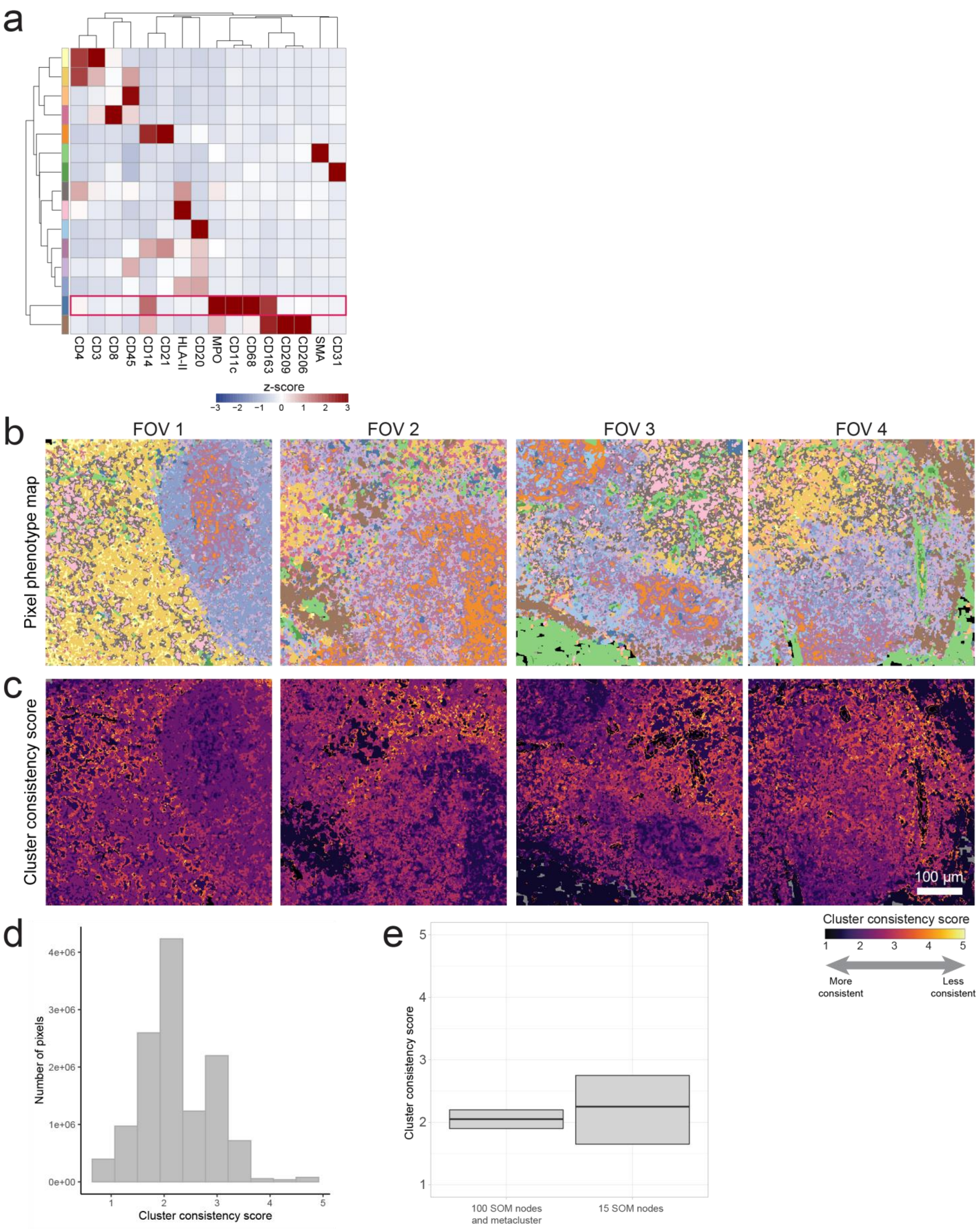

(A) Heatmap of mean marker expression of pixel cluster phenotypes where a SOM was used to cluster pixels directly into 15 clusters. The red box indicates an ambiguous pixel cluster with poor cluster definition. (B) Pixel phenotype maps for four representative FOVs (colored according to the heatmap in A). (C) The same FOVs in B colored according to cluster consistency score. (D) The distribution of cluster consistency score across all pixels in the dataset. (E) Comparison of the distribution of cluster consistency scores across all pixels in the dataset for metaclustering vs. directly clustering into 15 clusters.

### Supplementary Figure 10: Number of passes

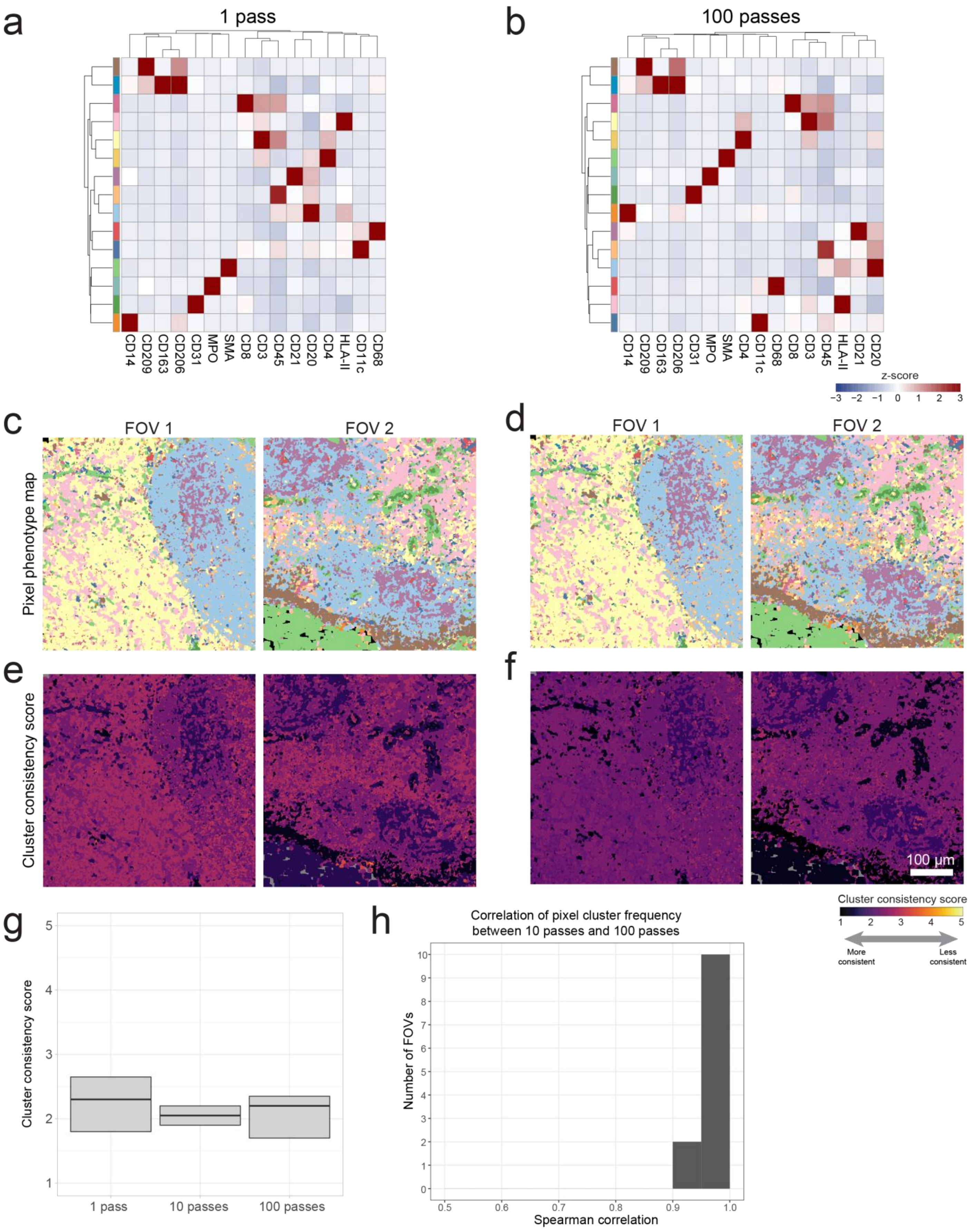

Heatmap of mean marker expression of pixel cluster phenotypes where the SOM was trained using 1 pass (A) or 100 passes (B). Pixel phenotype maps for two representative FOVs corresponding to 1 pass (C) or 10 passes (D). Pixel phenotype maps are colored according to A and B, respectively. The same FOVs in C and D colored according to cluster consistency score for 1 pass (E) or 10 passes (F). (G) Comparison of the distribution of cluster consistency scores across all pixels in the dataset for different number of training passes through the SOM. (H) Spearman correlation of pixel cluster frequency between 10 passes and 100 passes.

**Supplementary Figure 11: Subset pixels (decidua dataset)**

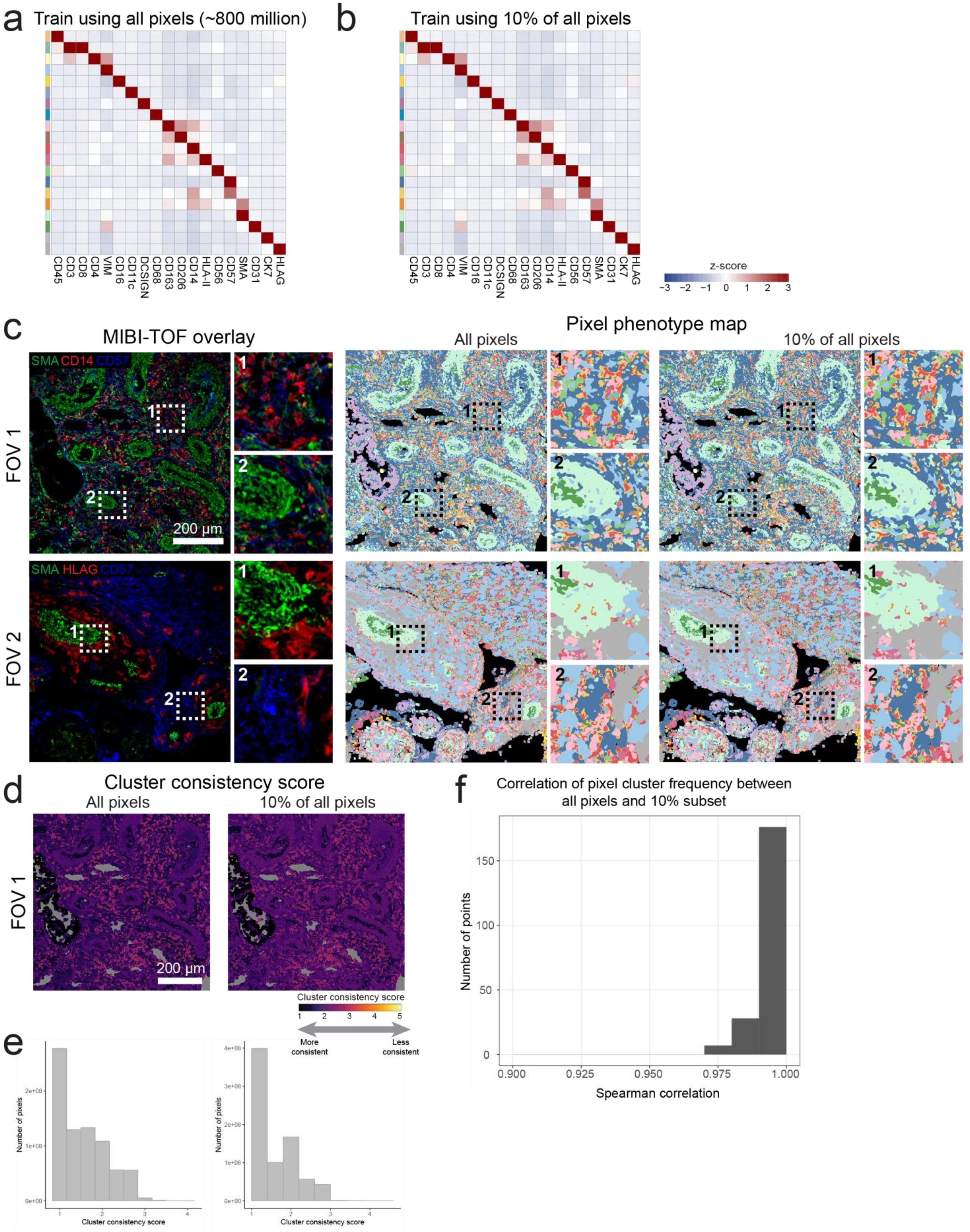

Heatmap of mean marker expression of pixel cluster phenotypes for a dataset of human decidua[33], where the SOM was trained using all pixels (A) or using a 10% subset of pixels (B). The full dataset contained a total of 766,440,566 pixels. (C) MIBI-TOF overlays (left) and pixel phenotype maps (right) for two representative FOVs. (D) FOV colored according to cluster consistency score for the SOM trained using all pixels (left) or a subset of pixels (right). (E) Comparison of the distribution of cluster consistency score across all pixels in the dataset for the SOM trained using all pixels (left) or a subset of pixels (right). (F) Spearman correlation of pixel cluster frequency between training using all pixels or a subset of pixels.

**Supplementary Figure 12: CyCIF whole slide tonsil dataset**

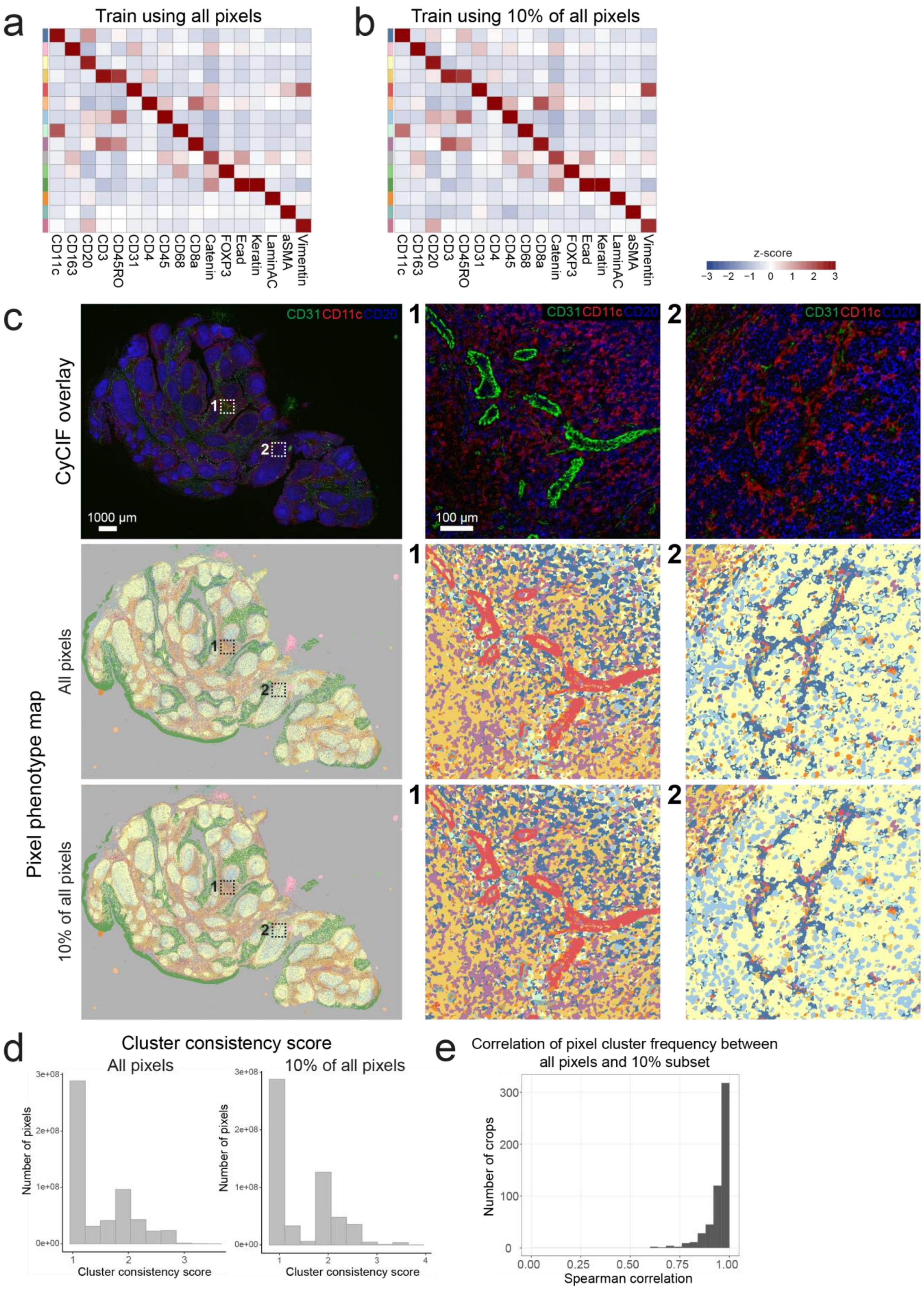

Heatmap of mean marker expression of pixel cluster phenotypes for a whole-slide CyCIF dataset of tonsil tissue[34], where the SOM was trained using all pixels (A) or using a 10% subset of pixels (B). The whole slide image was 27,299 x 20,045 pixels. (C) CyCIF overlays (top row) and pixel phenotype maps where the SOM was trained using all pixels (middle row) or using a 10% subset of pixels (bottom row). (D) Distribution of cluster consistency score across all pixels in the dataset for the SOM trained using all pixels (left) or a subset of pixels (right). (E) Spearman correlation of pixel cluster frequency between training using all pixels or a subset of pixels. To match the Spearman correlation calculation of the datasets with individual FOVs, 1024 x 1024 pixel crops were taken of the whole-slide image for the correlation calculation.

**Supplementary Figure 13: DCIS and TNBC pixel cluster profiles and TNBC quantification**

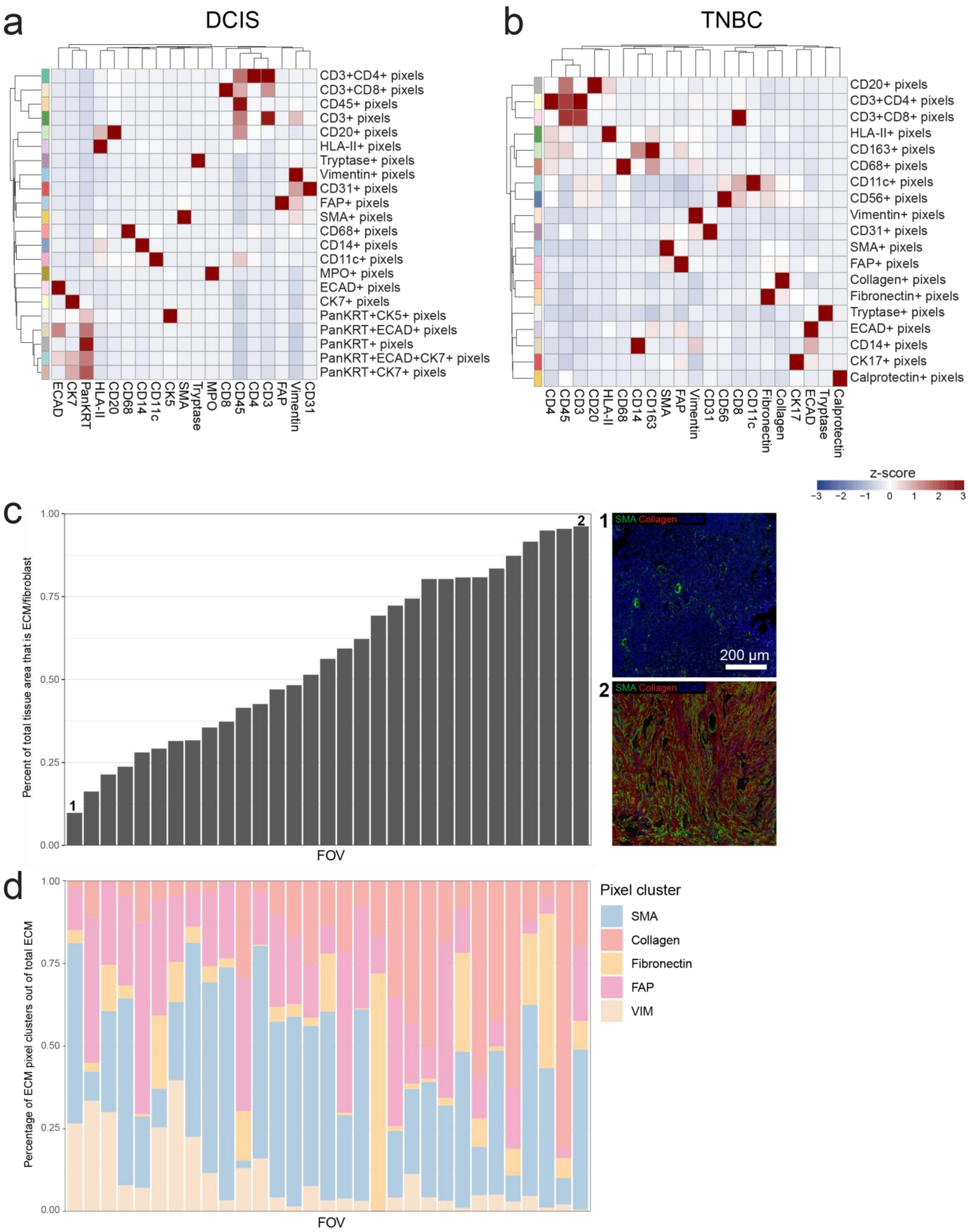

Pixel cluster expression profiles corresponding to the pixel clusters in Fig. 3b (DCIS) and Fig. 3c (TNBC) respectively. Expression values were z-scored for each marker. (C) For each FOV in the TNBC dataset, quantification of the percent of total tissue area that is comprised of pixel clusters of the ECM or fibroblast phenotypes. On the right, MIBI-TOF overlays of the FOVs with the lowest and highest amount of ECM/fibroblast. (D) Breakdown of the ECM/fibroblast pixel clusters for each FOV.

### Supplementary Figure 14: Reproducibility of pixel clustering

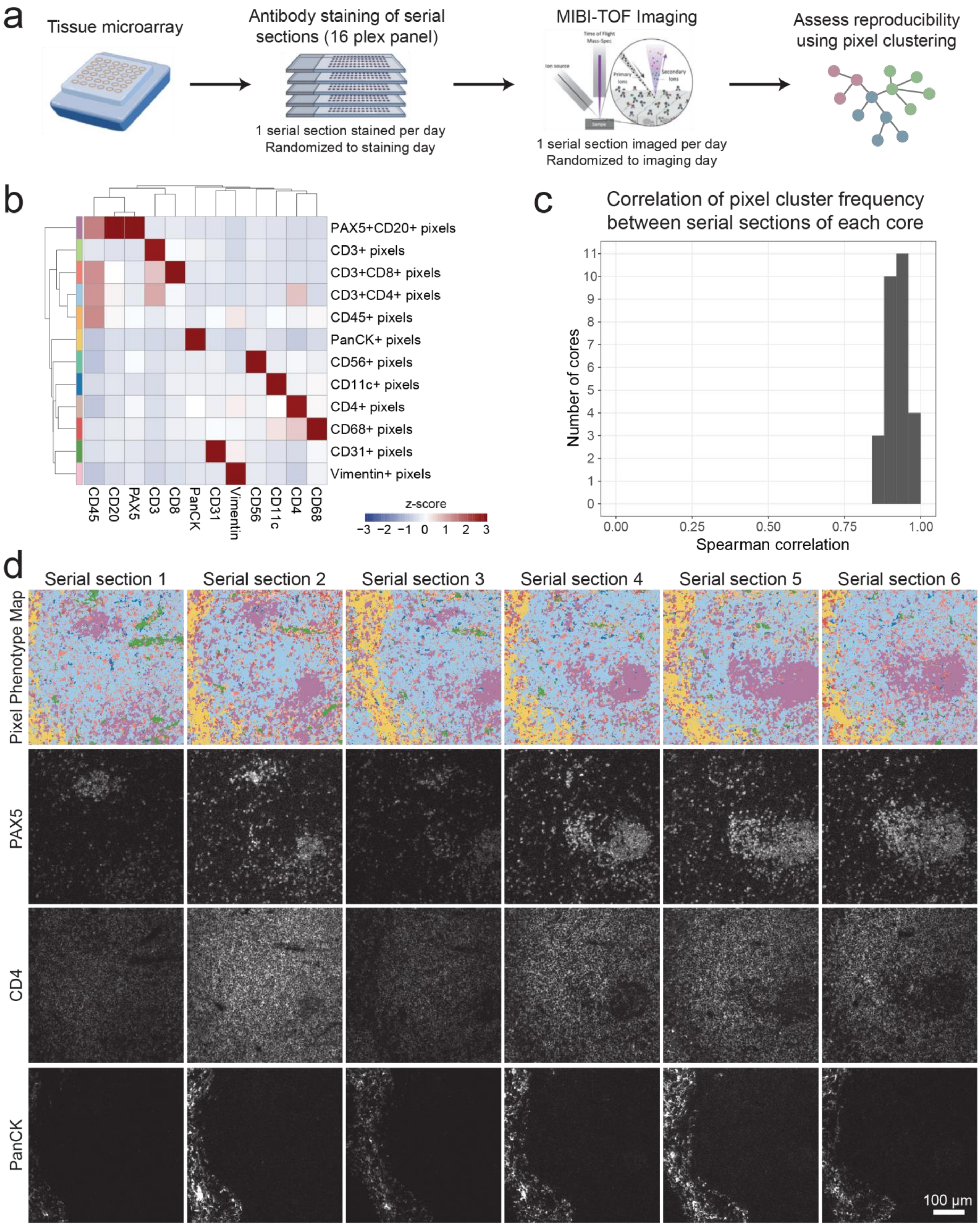

(A) A tissue microarray (TMA) comprised of various tissue types was serially sectioned, stained with a 16-plex panel, and imaged using MIBI-TOF.[35] The order that each serial section was stained and imaged was randomized. We then assessed the reproducibility of MIBI-TOF and pixel clustering by quantifying features between serial sections of the same TMA core. (B) Heatmap of pixel cluster phenotypes across the entire dataset. Expression values were z-scored for each marker. (C) The Spearman correlation between all serial sections of each TMA core using the frequency of pixel clusters in each FOV. (D) Example of pixel phenotype maps (colored according to the pixel clusters shown in B and single-channel images for six serial sections of the same tonsil tissue core. The single-channel images have the same maximum value.

**Supplementary Figure 15: Quantification of pixel cluster phenotypes in human hippocampus**

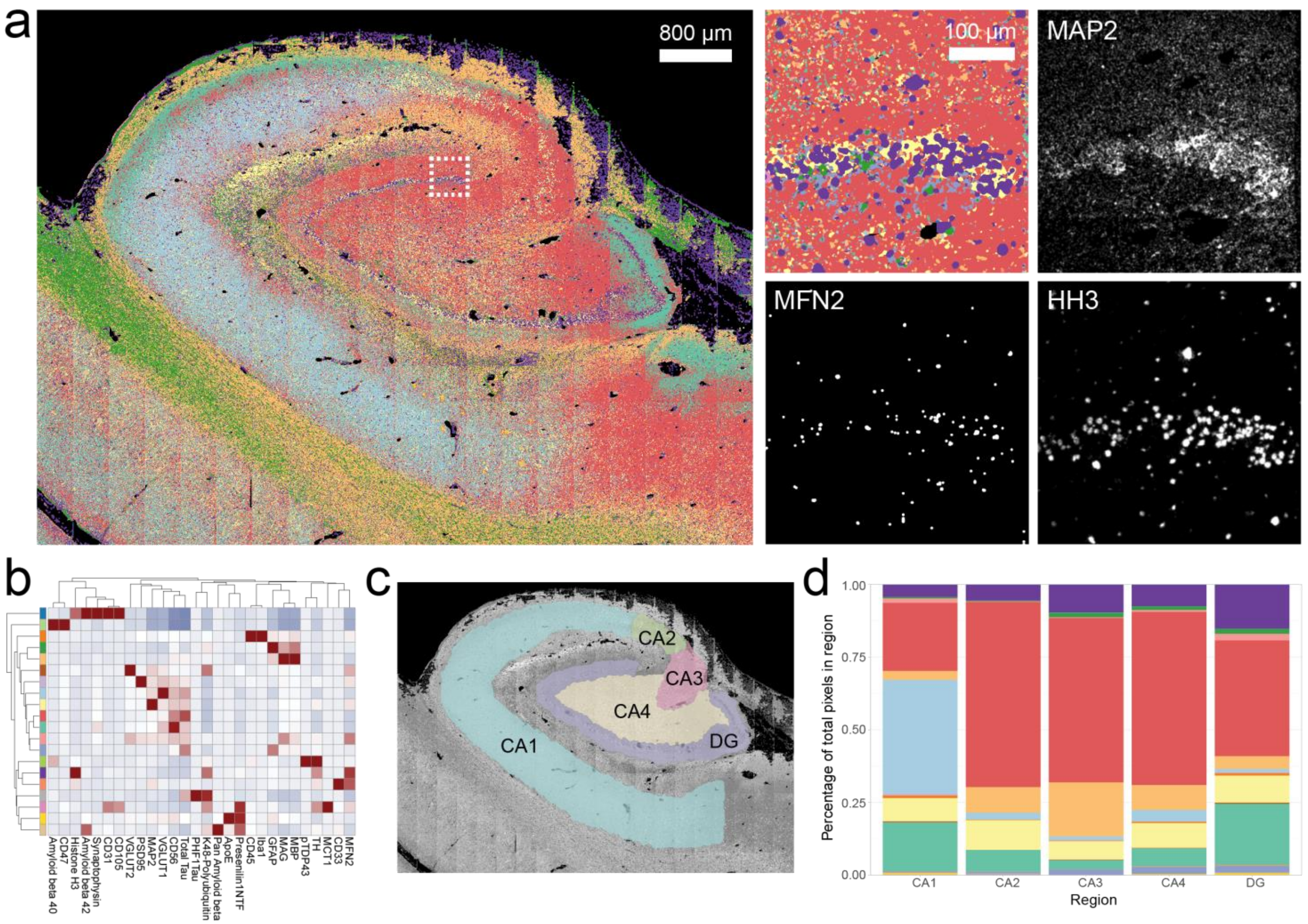


(A) Pixel phenotype map of MIBI-TOF image of cognitively normal human hippocampus tissue section[13]. 196 FOVs of 400 µm x 400 µm were captured by MIBI-TOF and tiled together. Insets (right) show local structure of the dentate gyrus, reflecting neuronal soma phenotypes as defined by MAP2, Histone H3, and MFN2 (mitofusion 2) expression. Total tiled MIBI-TOF image contained 63,642,954 non-zero pixels. (B) Heatmap of mean marker expression of pixel cluster phenotypes. Colors in the color bar correspond to the overlay in A. Proteins used for clustering include markers for microglia (CD45, Iba1) , astrocytes (GFAP), neurons (CD47, MAP2, TH, Tau, Synaptophysin, VGLUT1, VGLUT2, CD56, oligodendrocytes (MAG, MBP), vasculature (CD31, CD105, MCT1), proteopathy (Amyloid beta 40, Amyloid beta 42, PHF1Tau, Presenilin1NTF, pTDP43) and additional functional markers (Histone H3, MFN2, polyubiquitin 48, ApoE, CD33). (C) Hippocampus neuroanatomy as outline by expert neuropathologist. Dentate Gyrus (DG) and Cornu Ammonis (CA) regions 1-4 labelled. (D) Quantification of the pixel clusters belonging to each hippocampal region.

### Supplementary Figure 16: Cell clustering using pixel composition in Pixie

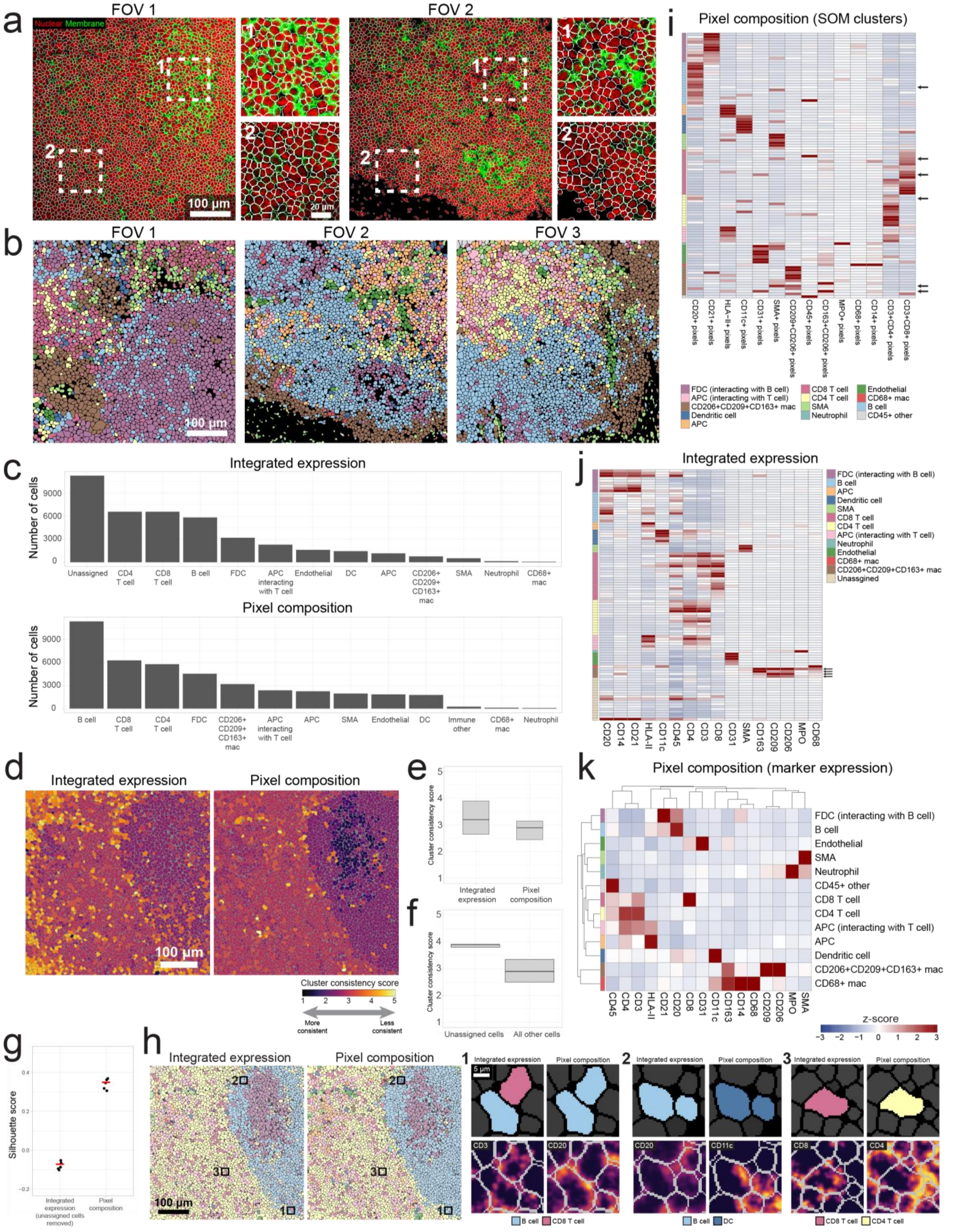

(A) Examples of segmentation quality. Images were segmented using the pre-trained Mesmer network (see Methods). We used histone H3 as the nuclear marker, and a combination of CD45, CD20, and HLA-II as the membrane marker. (B) Additional examples of cell phenotype types for representative FOVs where cells were clustered using pixel cluster composition. Colors in the cell phenotype maps correspond to the heatmap in Fig. 5c. (C) Total number of cells of each phenotype identified using integrated expression (top) or pixel composition (bottom). (D) The FOV shown in Fig. 5e colored according to cluster consistency score, for clustering using integrated expression (left) or pixel composition (right). (E) Comparison of cluster consistency score for cell clusters obtained using integrated expression or pixel composition. (F) For clustering using integrated expression, comparison of cluster consistency score for cells assigned to the unassigned group versus all other phenotypes. (G) Silhouette score comparison between cell clusters obtained using integrated expression, where cells in the "Unassigned" cluster were removed, and cell clusters obtained using pixel composition. (H) Cell phenotype maps of the FOV shown in Fig. 5e (left) and examples where the clustering was ambiguous or incorrect. (I) Heatmap of the 100 SOM clusters, clustered using pixel composition, grouped according to their final annotation. Arrows on the right correspond to clusters that had some ambiguous expression patterns that were manually inspected. (J) Heatmap of the 100 SOM clusters, clustered using integrated expression, grouped according to their final annotation. The arrows correspond to the CD206+ CD209+ CD163+ cluster, showing that all the individual clusters expressed the three markers with a z-score > 0. (K) Heatmap of marker expression for the cell phenotypes found using pixel composition. Marker expression was found by multiplying the number of each pixel cluster in each cell by the pixel cluster expression profile, then averaging across cells in the cluster.

**Supplementary Figure 17: Cell clustering using integrated expression from pre-processed pixel data**

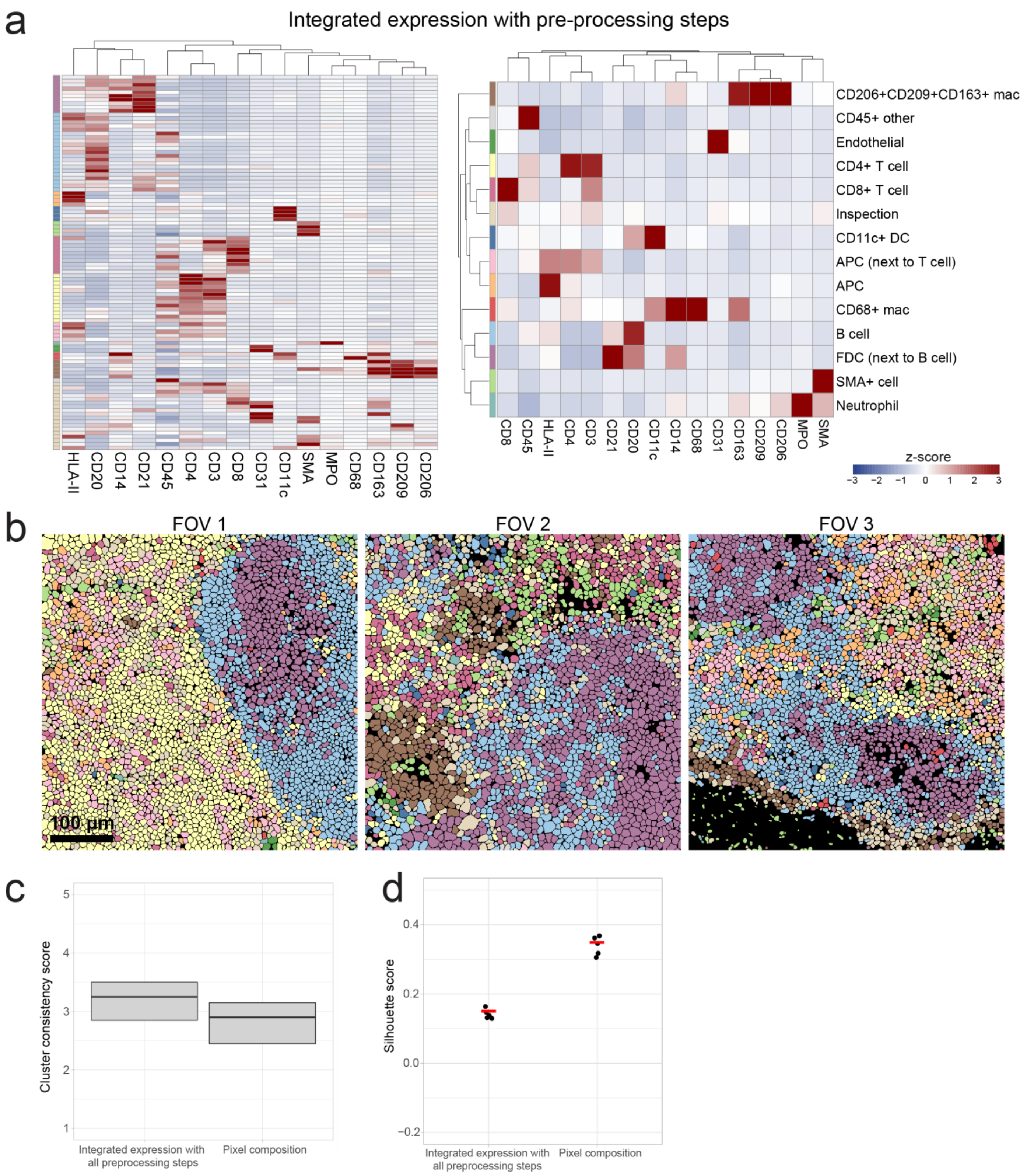


(A) Heatmap of the 100 SOM clusters (left) and annotated pixel cluster phenotypes (right) clustered using integrated expression, where the images were pre-processed as described for pixel clustering (i.e. Gaussian blur, pixel normalization, 99.9% normalization). (B) Cell phenotype maps for representative FOVs. Colors correspond to the heatmaps in A. (C)

Comparison of cluster consistency score between clustering using integrated expression of pre-processed pixel data or clustering using pixel composition. (D) Comparison of Silhouette score between clustering using integrated expression of pre-processed pixel data or clustering using pixel composition.

**Supplementary Figure 18: Cell clustering using segmentation from Ilastik/CellProfiler**

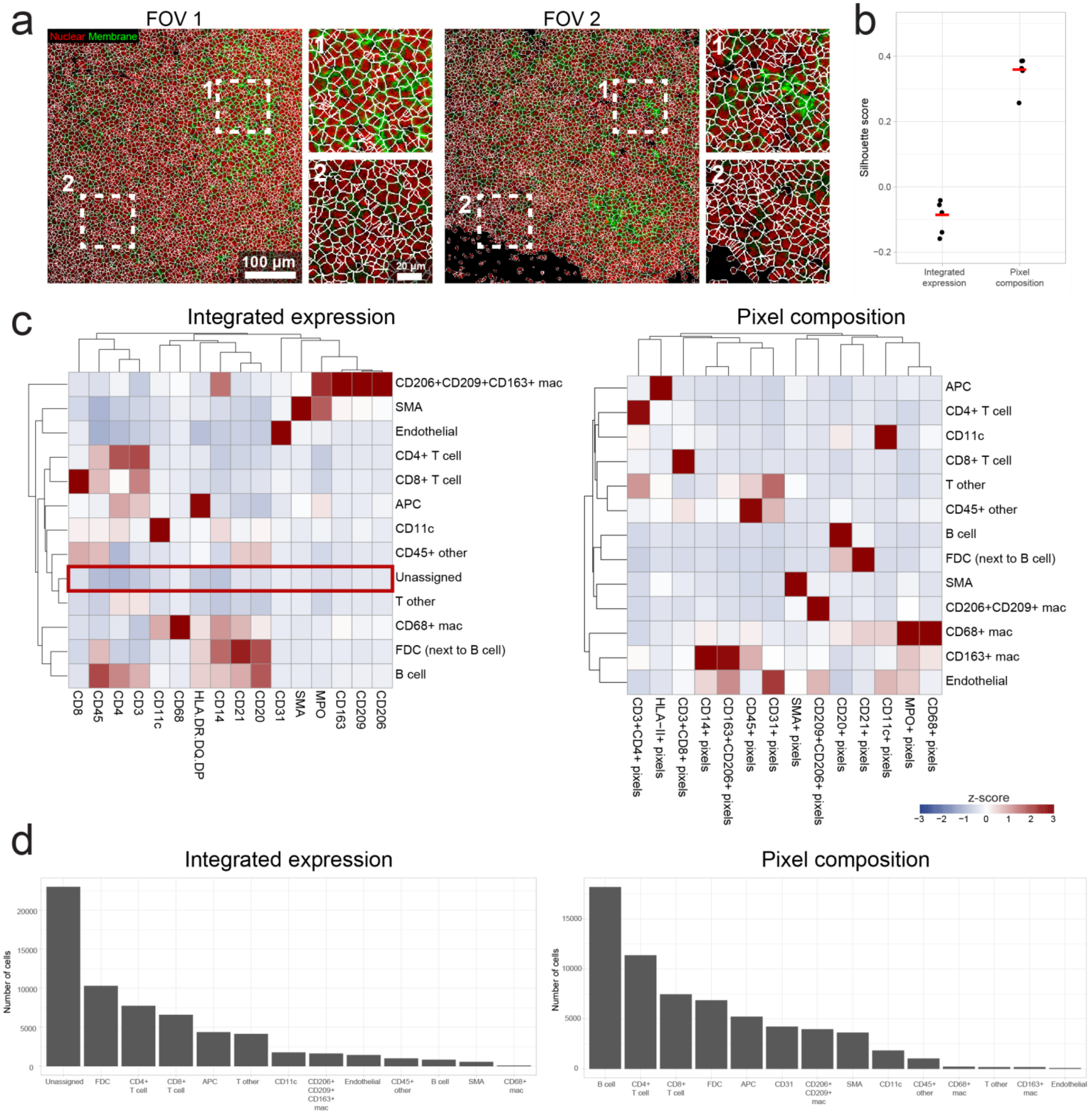


(A) Images showing segmentation performance using the Ilastik and CellProfiler segmentation pipeline.[51] We used histone H3 as the nuclear marker, and a combination of CD45, CD20, and HLA-II as the membrane marker. (B) Comparison of Silhouette score between pixel clustering using integrated expression or pixel composition, using the Ilastik/CellProfiler segmentation masks. (C) Heatmap of mean marker expression of cell cluster phenotypes using integrated expression (left) or pixel composition (right). (D) Total number of cells of each phenotype identified using integrated expression (left) or pixel composition (right).

### Supplementary Figure 19: Comparison of cell clustering using manually labeled dataset

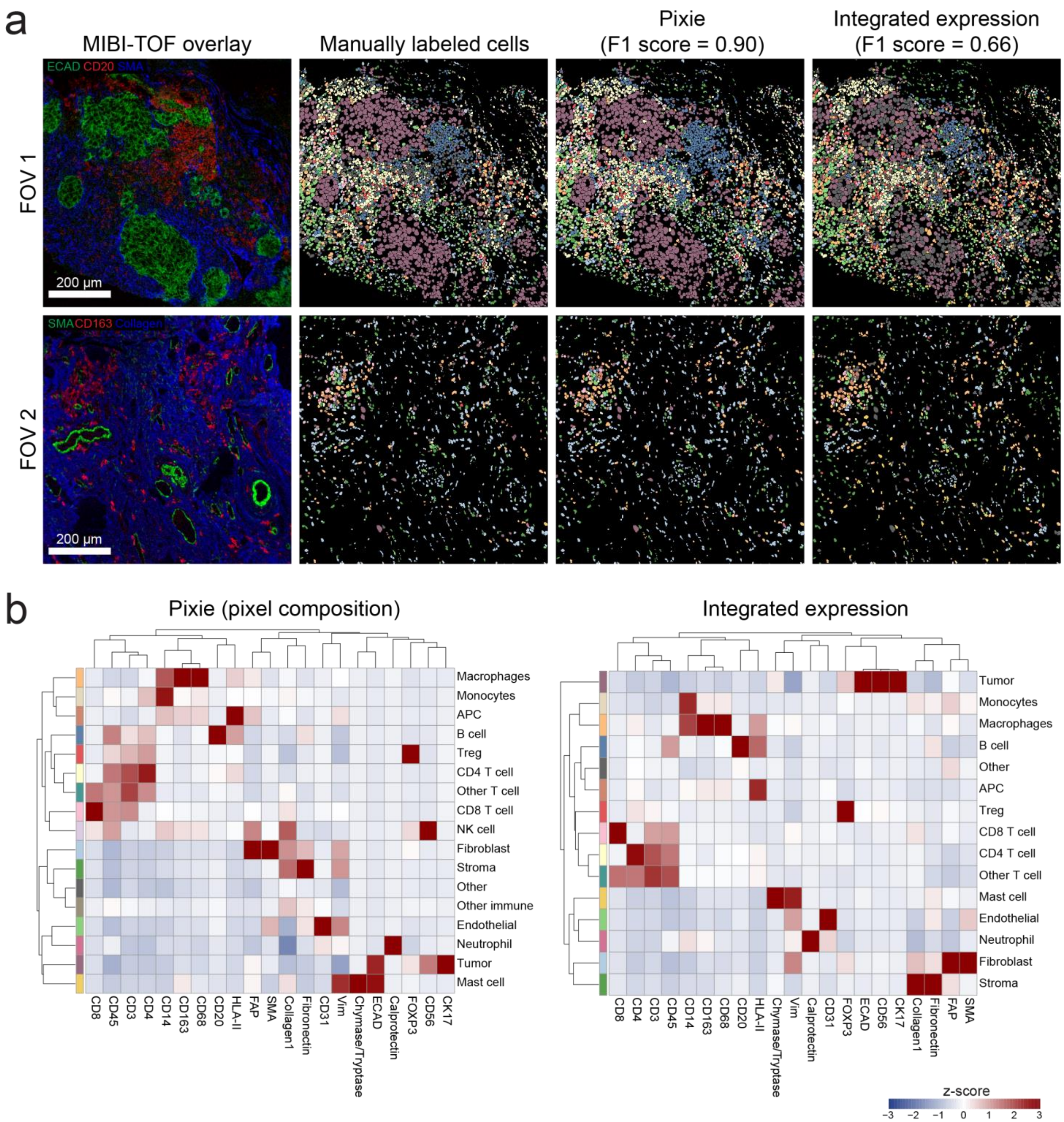


(A) Individual cells were manually annotated in MIBI-TOF images and used to assess cell clustering in Pixie. MIBI-TOF overlay (left) and cell phenotype maps (right) where cells are colored by their human-labeled phenotypes, where cells were clustered using pixel composition in Pixie, and where cells were clustered using integrated expression. (B) Heatmap of mean marker expression of the cell phenotypes identified using pixel composition (left) or integrated expression (right). The colors in the color bar correspond to the cell phenotype maps in A.

**Supplementary Figure 20: Comparison of Pixie with Otsu thresholding**

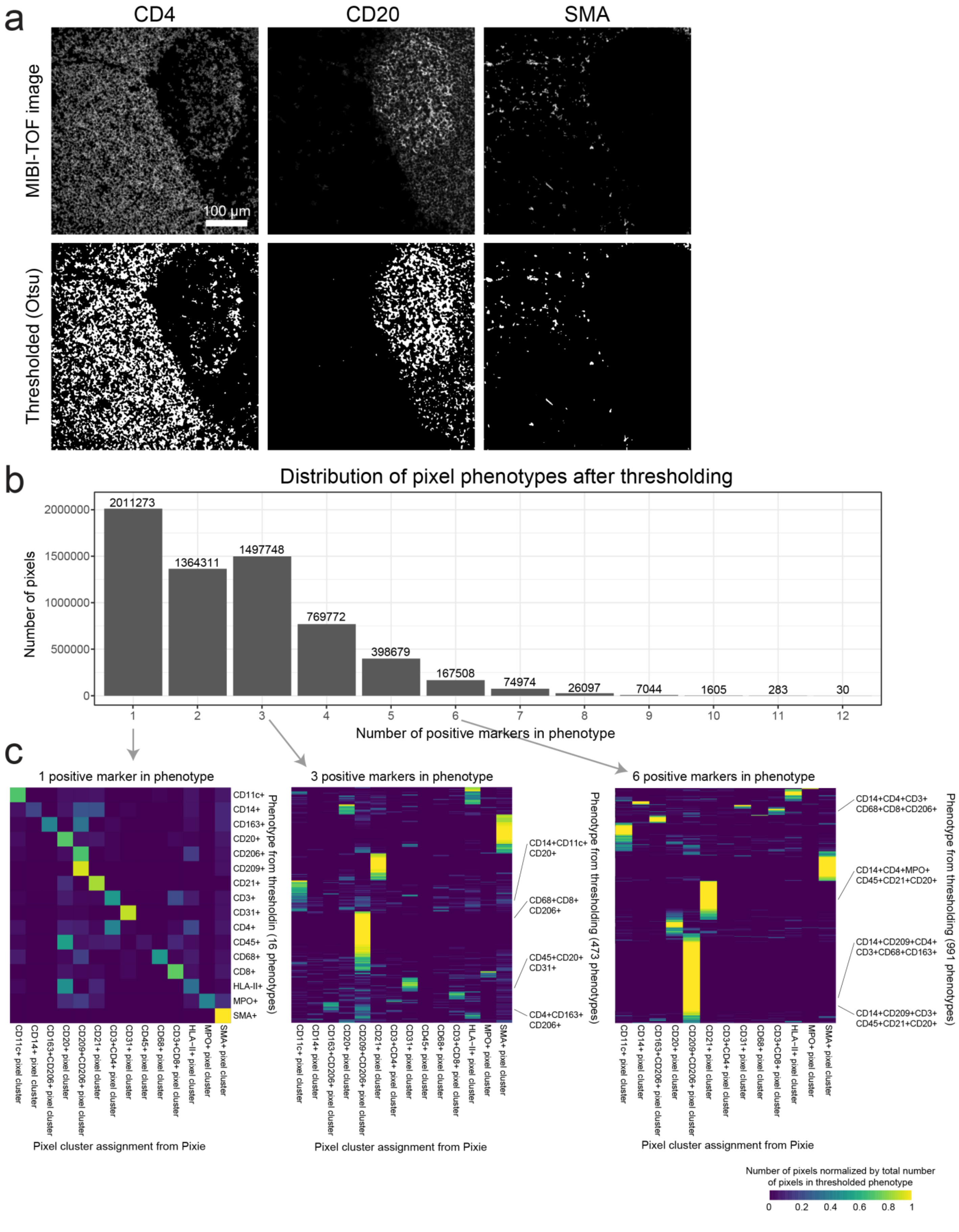

(A) Single-channel MIBI-TOF images (top row) were thresholded using Otsu's method to determine marker positivity. The thresholded images are shown in the bottom row (positive pixels are in white). Representative markers are shown. (B) For each pixel, we determined the number of markers that were called as positive using Otsu's method. Here, we are showing the distribution of the number of positive markers per pixel for the entire dataset. (C) Three representative examples showing the breakdown of the Otsu thresholded data (y axis) compared to the Pixie assignment (x axis). The heatmaps show the number of pixels normalized by the total number of pixels in the thresholded phenotype. For pixels that only contained 1 positive marker, there were 16 total phenotypes (i.e. the 16 markers included). For pixels that contained 3 positive markers, there were 473 total combinations, and for pixels that contained 6 positive markers, there were 991 total combinations.

**Supplementary Figure 21: Reproducibility of cell clustering using pixel cluster composition**

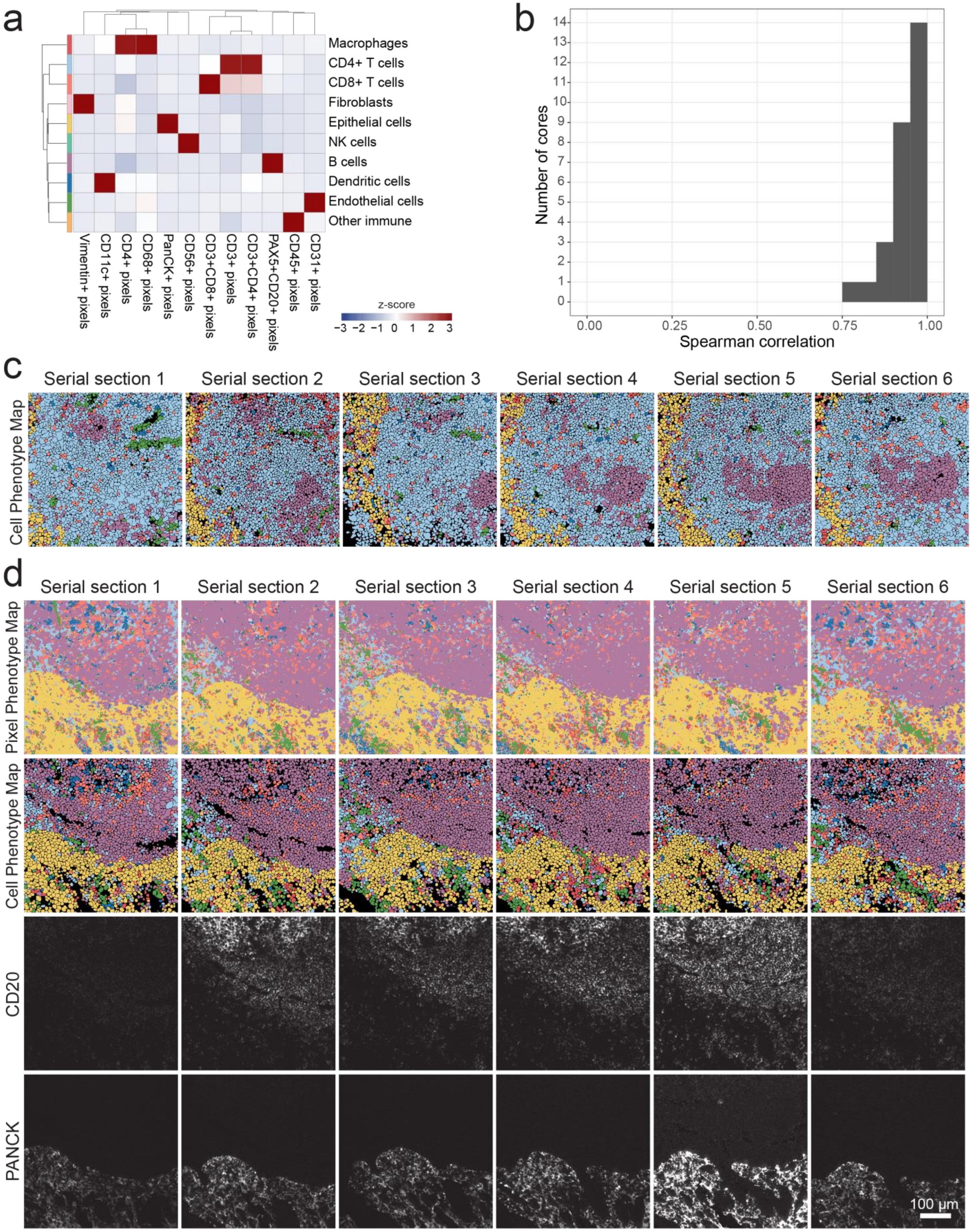

(A) Heatmap of cell phenotypes for the MIBI-TOF dataset in the reproducibility study shown in Supplementary Fig. 14.[35] (B) The Spearman correlation between all serial sections of each TMA core using the frequency of cell clusters in each FOV. (C) Cell phenotype maps (colored according to the cell phenotypes shown in A for the tonsil tissue core shown in Supplementary Fig. 14d. (D) Additional example of pixel phenotype maps colored according to the pixel clusters shown in Supplementary Fig. 14 (top row), cell phenotype maps colored according to the cell clusters shown in A (second row), and single-channel images (third and fourth rows) for six serial sections of the same tonsil tissue core. The single-channel images have the same maximum value.

### Supplementary Figure 22: Runtime analysis

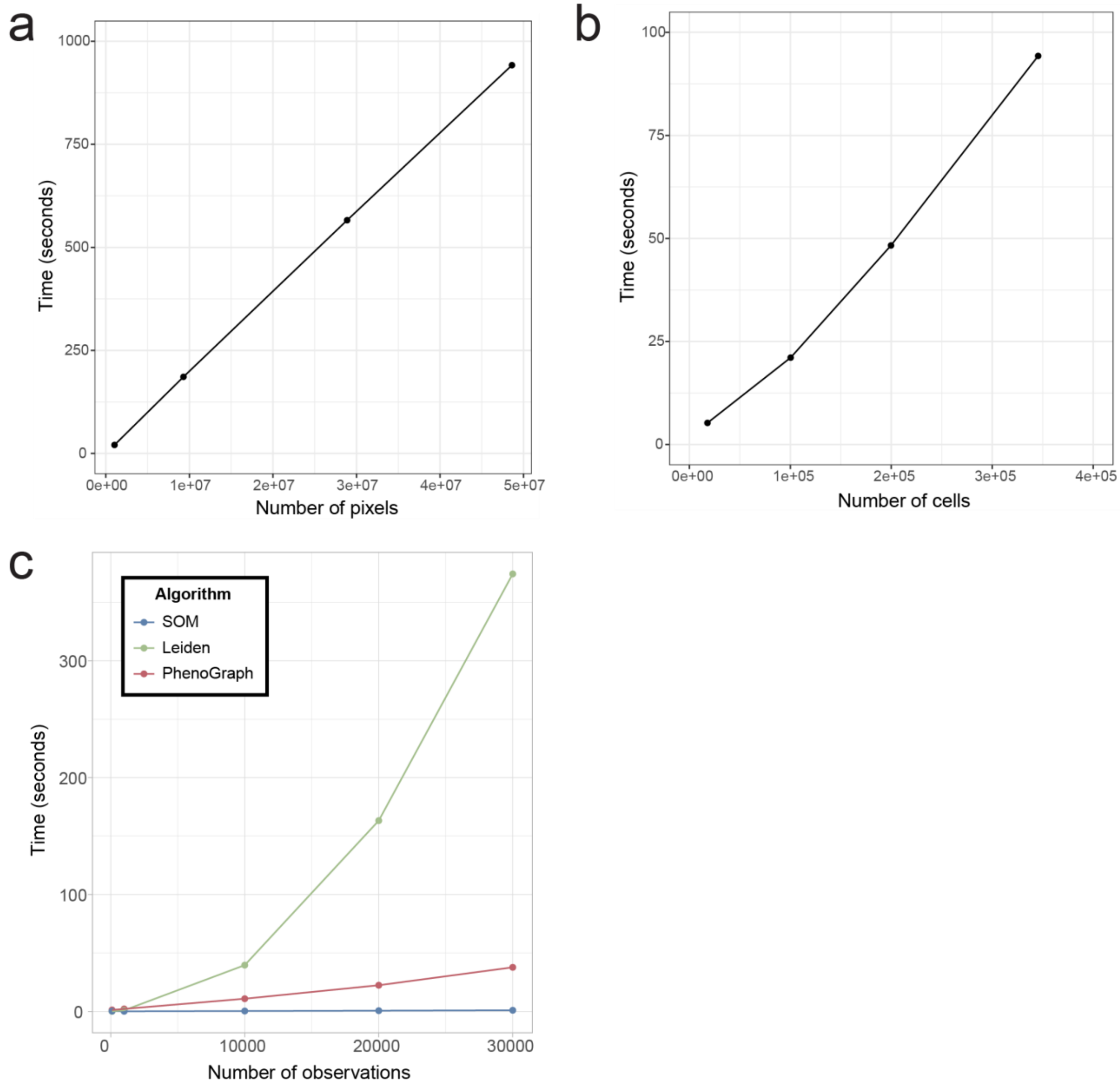


(A) Total Pixie runtime including all pre-processing steps and clustering for pixel clustering. (B) Total Pixie runtime including all pre-processing steps and clustering for cell clustering. (C) Runtime comparison between SOM (implemented in FlowSOM), Leiden (implemented in Seurat), and PhenoGraph (implemented in Rphenograph) clustering algorithms. Runtime comparison was performed on a Google Cloud Compute Engine instance with 16 vCPU and 128 GB of memory.
